# Genome-wide mapping of helicase-generated ssDNA reveals Hrq1 activity at RNA polymerase III-transcribed genes

**DOI:** 10.64898/2026.08.13.744683

**Authors:** Shaili Regmi, Noof Alsulaiti, Daniel Darling, Anastasiya Bolgova, Bertrand Theulot, Spencer J. Gray, Matthew L. Bochman, Duncan J. Smith

## Abstract

DNA helicases preserve genome stability by unwinding DNA during replication, repair, recombination, and transcription, yet their sites of action *in vivo* remain difficult to define. Here, we describe a sequencing-based strategy to map helicase activity genome-wide by coupling helicases to the single-stranded DNA-specific activation-induced cytidine deaminase (AID). Deamination of cytosines exposed during helicase-mediated DNA unwinding generates strand-specific mutational footprints that can be detected by whole-genome sequencing at near-nucleotide resolution. Using the *Saccharomyces cerevisiae* RecQ4-family helicase Hrq1, a functional homolog of human RECQL4, we generated the first genome-wide map of Hrq1 activity. Hrq1-dependent deaminations were highly enriched at RNA polymerase III (RNAPIII)-transcribed genes, particularly tRNA genes, where they occurred predominantly on the transcriptional template strand. This localization was reproducible using both overexpressed Hrq1-AID fusions and an inducible dimerization system that recruited AID to endogenously expressed Hrq1, and it was markedly reduced by helicase-inactivating mutation, indicating that active DNA unwinding underlies the observed signal. Hrq1 associated with nearly all tRNA genes irrespective of transcription level, replication orientation, or proximity to transposable elements, yet deletion or overexpression of Hrq1 did not detectably alter pre-tRNA abundance or RNA polymerase III recycling under the conditions tested. Application of the same approach to the PIF1-family helicase Rrm3 recovered its established enrichment at a subset of highly transcribed, head-on tRNA genes, validating the method. Together, these findings establish AID-mediated mutational footprinting as a general approach for mapping helicase activity *in vivo* and reveal an unexpected, widespread association of the RecQ4-family helicase Hrq1 with RNAPIII-transcribed genes.

## Introduction

Helicases are nucleic acid–dependent ATPases that translocate along DNA or RNA and use the energy derived from ATP hydrolysis to unwind or remodel nucleic acid structures (1, 2). DNA helicases use this biochemical activity to support genome integrity, participating in pathways such as DNA replication, recombination, and repair, transcription, telomere maintenance, and the resolution of secondary structures like R-loops and G-quadruplexes (G4s) (1). Defects in these processes can lead to genomic instability and disease. Thus, DNA helicases are highly conserved and essential for maintaining genome stability (3–5).

The RecQ helicase family represents one such group of evolutionarily conserved enzymes, found from bacteria to humans (6, 7). These 3’ to 5’ Superfamily 2 (SF2) helicases are often called the “guardians of the genome” because of their central roles in maintaining genome integrity. The human genome encodes five RecQ helicases (RECQL1, RECQL2/WRN, RECQL3/BLM, RECQL4, and RECQL5). Mutations in RECQL1, BLM, WRN, and RECQL4 cause inherited genetic diseases characterized by genomic instability, developmental abnormalities, features of premature aging, and/or predispositions to cancer (7, 8).

Mutations in RECQL4 cause three different autosomal recessive diseases (Rothmund-Thomson syndrome (9), Baller-Gerold syndrome (10), and RAPADILINO (11)), and the Catalogue of Somatic Mutations in Cancer (12) categorizes *RECQL4* as both a tumor suppressor and oncogene due to its known roles in supporting genome stability and the dozens of *recql4* mutations found in cancers (13–15). However, RECQL4 remains one of the least-understood human RecQ helicases because it is technically challenging to work with. Although the C-terminal two-thirds of the protein contain a conserved RecQ helicase core domain where the disease alleles are found (16), its large N-terminal domain resembles the *Saccharomyces cerevisiae* Sld2 replication initiation factor at the primary sequence level (17–20). It is unclear if RECQL4 acts in a Sld2-like capacity in metazoans (21), but it is well established to have a role in replication initiation, which does not require helicase activity (17–20). As such, attempting to study disease-related mutations can have off-target effects on replication. Because RECQL4 is large (>130 kDa) and its Sld2-like domain is natively disordered (19), it is also difficult to generate recombinantly for *in vitro* experiments (22, 23).

To circumvent these challenges, we have established the *S. cerevisiae* Hrq1 helicase as an experimentally tractable functional homolog of the helicase portion of RECQL4 (23–35). Hrq1 is a biochemically amenable enzyme that shares a conserved RecQ4-family helicase core domain and structural, biochemical, and functional similarities with RECQL4. Unlike metazoan RECQL4, however, Hrq1 does not combine RecQ4-family helicase functions with a Sld2-like replication initiation module (16) because Sld2 is encoded as a separate protein in budding yeast (36). Thus, *S. cerevisiae* and Hrq1 represent a powerful experimental system to dissect the conserved functions of RecQ4-family helicases without perturbing the essential Sld2 pathway.

To date, we have reported conserved functions of Hrq1 and RECQL4 in DNA interstrand crosslink (ICL) repair (23, 31, 33, 35, 37, 38) and telomere maintenance (23–28, 33). Hrq1 also exhibits genetic synergism with Sgs1, the budding yeast homolog of the human BLM helicase (23–25, 34). At the biochemical level, Hrq1 and RECQL4 exhibit related DNA binding, ATPase, and helicase activities, including the ability to unwind G4 structures (23, 30, 31, 33). Together, these findings establish Hrq1 as a useful model for investigating conserved RecQ4-family functions.

Having used Hrq1 to investigate functions attributed to RECQL4, we next asked if we could use it as a discovery platform to identify previously unrecognized activities of RecQ4-family helicases. We reasoned that if we could identify sites of Hrq1 activity genome wide, we could use the data to generate testable hypotheses for why Hrq1 is active at those loci. For instance, despite not having a Sld2-like domain (16), if Hrq1 localizes to origins of replication, then it likely functions in replication initiation. Initially, we attempted to do this via ChIP-seq and ChEC-seq, but Hrq1-3xFLAG and Hrq1-MNase (respectively) were refractory to these techniques. Therefore, we developed a novel method in which we tagged Hrq1 with a genome editing enzyme, expressed the fusion *in vivo*, and conducted whole genome sequencing to identify the mutational fingerprint of the chimeric enzyme. This approach yielded highly repeatable, stranded, and nucleotide-resolution data, enabling us to generate a genome-wide Hrq1 activity map. Unexpectedly, we found that Hrq1 is highly enriched at genes transcribed by RNA polymerase III (RNAPIII), and the consequences of this as they pertain to RECQL4 are discussed below.

## Materials and Methods

### Reagents

The following enzymes were used: T4 Polynucleotide Kinase, 3ʹ-phosphatase minus (New England Biolabs; Ipswitch, MA, USA; M0236L) and TURBO DNase (Invitrogen; Carlsbad, CA, USA; AM2238). The following kits were used: Illumina DNA Prep Kit (San Diego, CA, USA; 20049005), IDT for Illumina Nextera DNA UD Indexes (Coralville, IA, USA; 20026934), RNA Clean & Concentrator-5 (Zymo Research; Irvine, CA, USA; R1015), and iTaq Universal SYBR Green One-Step Kit (Bio-Rad; Hercules, CA, USA; 725150). The following commercial instruments were used: Illumina NovaSeq 6000 (San Diego, CA, USA; 20012850), SequaGel UreaGel System (National Diagnostics; Atlanta, GA, USA; EC-833), Trans-Blot^®^ SD Semi-Dry Electrophoretic Transfer Cell (Bio-Rad; Hercules, CA, USA;170-3940), Amersham Biosciences UVC 500 Crosslinker (Piscataway, NJ, USA; 80-6222-31), and a CFX96 Real-Time PCR Detection system (Bio-Rad; Hercules, CA, USA; 184-5096). The following chemicals were used: Rapamycin (bioWORLD; Dublin, OH, USA; 41810000), hydroxyurea (HU) (Sigma-Aldrich; St. Louis, MO, USA; H8627), and [γ-^32^P]ATP (Revvity; Waltham, MA, USA; BLU502A100UC).

### Biological Resources

All *S. cerevisiae* strains used in this study are derivatives of W303 *RAD5+* (*ade2-1 can1-100 his3-11, 15 leu2-3, 112 trp1-1 ura3-1 RAD5+)* or of S288C (*SUC2 gal2 mal2 mel flo1 flo8-1 hap1 ho bio1 bio6*) and are listed in Supplementary Table 1. Strains were generated by LiAc/SS Carrier DNA/PEG transformation or by mating, sporulation, and tetrad dissection, as previously described (39). Cells were grown in yeast extract, peptone, dextrose (YEPD, 2% glucose) medium at 30℃ unless otherwise stated.

### Induction conditions for activation-induced cytidine deaminase (AID)-based mapping

*hrq1Δ* and *hrq1Δ ung1Δ* strains (with *pGAL-HRQ1-AID* or control plasmids) were grown from an optical density at 600 nm (OD_600_) of 0.2 to 0.3 in -URA + 2% glucose at 30°C (lag-phase recovery), washed twice, and shifted to -URA + 1.5% raffinose + 0.5% galactose for induction (4 doublings). Rapamycin-resistant (*tor1-1 fpr1Δ*) *HELICASE-FRB, pGAL-AID-FKBP12 ung1Δ* strains were grown from OD_600_ 0.2 to 0.3-0.5 in YEPD (2% glucose) + adenine (0.12 mg/mL) at 30°C, washed twice, and shifted to YEPGal (2% galactose) + adenine + 2 µg/mL rapamycin for induction/dimerization. Samples were collected at 1 day (5 doublings) and after back-dilution of the 1-day culture to OD_600_ 0.2 followed by 1 more day of growth (2-day sample, 10 doublings). Controls were grown in parallel in glucose + rapamycin or galactose without rapamycin; 10 mM HU was added where specified. All samples were pelleted and frozen at -20°C before DNA extraction and sequencing.

### DNA extraction and sequencing

DNA was extracted as previously described (40). Sequencing libraries were prepared using an Illumina DNA Prep Kit (20049005) and barcoded using IDT for Illumina Nextera DNA UD Indexes (20026934) according to the manufacturer’s instructions (Illumina, Inc.). Libraries were sequenced on an Illumina NovaSeq 6000 using an SP flow cell with 150 bp paired-end reads (300 cycle v1.5 kit). Using a custom nextflow pipeline, GENEFLOW (41), reads were basecalled (Picard IlluminaBasecallsToFastq v2.23.8 with APPLY_EAMSS_FILTER set to false) and demultiplexed (Pheniqs v2.1.0) (42).

### Custom reference genome generation

A custom W303-1A *RAD5+* reference genome was sequenced in-house. Genomic DNA was extracted as previously described (40), and Nanopore sequencing libraries were prepared per the manufacturer’s instructions (Oxford Nanopore Technologies). Duplex reads were basecalled from POD5 files using Dorado v0.5.1 (super-accuracy duplex mode, GPU-accelerated), retaining only duplex-classified reads with Phred quality score (Q) >= 20. Filtered reads were aligned to the S288C reference (sacCer3) using minimap2 v2.26-r1175 (map-ont preset) and processed with SAMtools v1.19; mitochondrial reads were excluded by extracting nuclear read IDs with SAMtools and seqtk v1.2-r94, and the nuclear read set was downsampled to 100× coverage (Rasusa v2.1, assuming a 12.4 Mb genome). *De novo* assembly was performed with Canu v2.2 (-nanopore-raw), and contigs were scaffolded against S288C using Ragout v2.3 (S288C as both reference and naming source). Assembly completeness was 99.3% by BUSCO, comparable to S288C. The scaffolded genome was annotated using LRSDAY v1.7.2. A separate mitochondrial assembly was generated from the mitochondrial read set using Flye v2.9.6-b1802.

### Sequence alignment and variant calling

Paired reads were aligned to the custom W303-1A *RAD5+* reference genome using BWA-MEM v0.7.17 with default parameters. BAM files were coordinate-sorted and indexed using SAMtools v1.14. Alignments bearing secondary (XA:Z:) or supplementary/chimeric (SA:Z:) tags were removed, and reads with mapping quality (MAPQ) >= 30, excluding unmapped/secondary/supplementary/duplicate flags (flag:3332) and requiring the properly paired flag (flag:2), were retained. PCR duplicates were identified and removed using Picard MarkDuplicates v2.27.5 with the REMOVE_DUPLICATES=true option. Deduplicated BAM files were indexed and pileup files generated using SAMtools mpileup against the custom reference genome. Single-nucleotide polymorphisms (SNPs) were called from the pileups using VarScan v2.4.2 (*p*-value threshold 0.95, minimum variant allele frequency 0.001, VCF output, other parameters default).

### Computational analyses and data source

All analyses and graphs were generated using custom R scripts (available upon request). Gene and transposable element coordinates were drawn from the annotated custom W303-1A *RAD5+* reference. ARS consensus sequence (ACS) sites were obtained from Eaton et al., 2010 (43) and remapped onto the custom reference. G4 sites were predicted on the custom reference using G4Hunter (44). tRNA intron information was obtained from GtRNAdb (sacCer3, Apr 2011) (45) and transferred by gene name matching. Replication orientation (head-on *vs.* co-directional) and leading/lagging strand assignment were derived from OK-seq data (46, 47) remapped onto the custom reference. tRNA transcription levels were inferred from RNAPIII occupancy (NET-seq read counts normalized to total reads) (48), matched to the custom reference by gene name.

### Cell growth and RNA extraction for northern blots and RT-qPCR

For northern blots, wild-type and *hrq1Δ* cells were grown to exponential phase at 30°C in YEPD. For Tfc1 depletion, strains were grown similarly and treated with 1 µg/mL rapamycin (via Anchor Away) (49), with cells harvested at 0, 20, 40, 60, and 90 min post-treatment. For RT-qPCR, wild-type, *hrq1Δ*, *maf1Δ*, and *hrq1Δ maf1Δ* strains were grown to exponential phase in YEPD at 16, 30, and 37°C. For overexpression assays, *hrq1Δ* strains carrying empty vector, *pGAL-HRQ1*, or *pGAL-HRQ1-KA* were grown to exponential phase in -URA + 2% glucose at 30°C, washed thrice, switched to -URA + 2% galactose, and collected hourly from 0-4 h post-induction. For all conditions, cells were harvested by centrifugation at 4°C, washed with 1X PBS, and pellets stored at -80°C prior to RNA extraction using a TRIzol-based method for yeast (50).

### Northern blotting

RNA (10 µg) was denatured in formamide buffer (95°C, 10 min; 95% formamide, 20 mM EDTA, 0.05% bromophenol blue, and 0.05% xylene cyanol) and resolved on a pre-run 10% polyacrylamide-urea gel (25 W, 1 h; SequaGel UreaGel System, National Diagnostics, EC-833), then stained with SYBR Gold (Invitrogen, S11494). RNA was semi-dry transferred to a Zeta-Probe membrane (Bio-Rad, 162-0159) using a Semi-Dry apparatus (Bio-Rad, 170-3940) and UV-crosslinked (1200 × 100 µJ/cm², 5 min; Amersham Biosciences UVC 500 Crosslinker). Transfer was confirmed by methylene blue staining (0.3 M sodium acetate, pH 5.5, 0.02% methylene blue). Membranes were blocked (37°C, overnight) with 125 µg/mL fish sperm DNA in 6× SSPE, 0.1% SDS, 2× Denhardt’s, and hybridized overnight under the same conditions with probes labelled with [γ-^32^P]ATP (Revvity, BLU502A100UC; T4 Polynucleotide Kinase, 3ʹ-phosphatase minus, NEB, M0236L) against SNR190 and the tL(CAA) intron. Membranes were washed (6× SSPE, 0.1% SDS), exposed to phosphorimager screens, and imaged on an Amersham Typhoon biomolecular imager. The probe for SNR190 was 5ʹ-GTCATGGTCGAATCGGACGAGG-3ʹ, and that for the tL(CAA) intron was 5ʹ-TATTCCCACAGTTAACTGCGGTCAAGATATTT-3ʹ.

### RT-qPCR

Total RNA was DNase-treated (TURBO DNase; Invitrogen, AM2238) and purified (RNA Clean & Concentrator-5; Zymo Research, R1015). 100 ng RNA per sample was analyzed by one-step RT-qPCR (iTaq Universal SYBR Green One-Step Kit; Bio-Rad, 725150; CFX96 system) using the primers 5ʹ-GGTTGTTTGGCCGAGCG-3ʹ and 5ʹ-CTTACGATACCTGAGTATTCCCACAG-3ʹ for the tL(CAA) pre-tRNA and 5ʹ-GTACTCTTCCGGTAGAACTACTGG-3ʹ and 5ʹ-CGATTCTCAAAATGGCGTGAGG-3ʹ for *ACT1* mRNA, each at 300 nM. Samples were run in technical triplicate with No-RT controls to confirm the absence of genomic DNA contamination. tL(CAA) pre-tRNA levels were normalized to *ACT1* mRNA and expressed as the log_2_-fold change relative to wild-type or empty vector control. A second biological replicate was performed independently.

### Serial dilution growth assays

Overnight cultures were grown in YEPD or -URA + 2% glucose, as appropriate, diluted to OD_600_ 0.65, and serially diluted 1:5. Dilutions were spotted onto the appropriate plates and grown at 30°C for 2 days unless otherwise stated.

### Statistical Analyses

Statistical methods and relevant details are provided in the appropriate sections of the main text and figure legends.

### Novel Programs, Software, Algorithms

Not Applicable.

### Web Sites/Data Base Referencing

Saccharomyces Genome Database (SGD) (51) and Genomic tRNA Database (GtRNAdb) (45).

## Results

### Direct AID fusion maps Hrq1 to RNAPIII-transcribed genes

Helicase unwinding of duplex DNA exposes single-stranded DNA (ssDNA), generating a substrate for ssDNA-modifying enzymes. We reasoned that targeting the ssDNA-specific cytidine deaminase AID to sites of helicase activity via direct fusion would introduce local deaminations and enable high-resolution mapping of helicase activity *in vivo*. AID is a member of the APOBEC (apolipoprotein B mRNA editing catalytic polypeptide-like) family that specifically deaminates cytosine residues to uracil within WRC motifs (W = A or T; R = A or G) on ssDNA (52). Deaminations result in C-to-T or G-to-A mutations following DNA replication and sequencing, depending on the affected strand. Mapping these mutation patterns across the genome should therefore provide a strand-specific, readout of helicase engagement with ssDNA *in vivo* (Fig. 1A).

**Figure 1.**
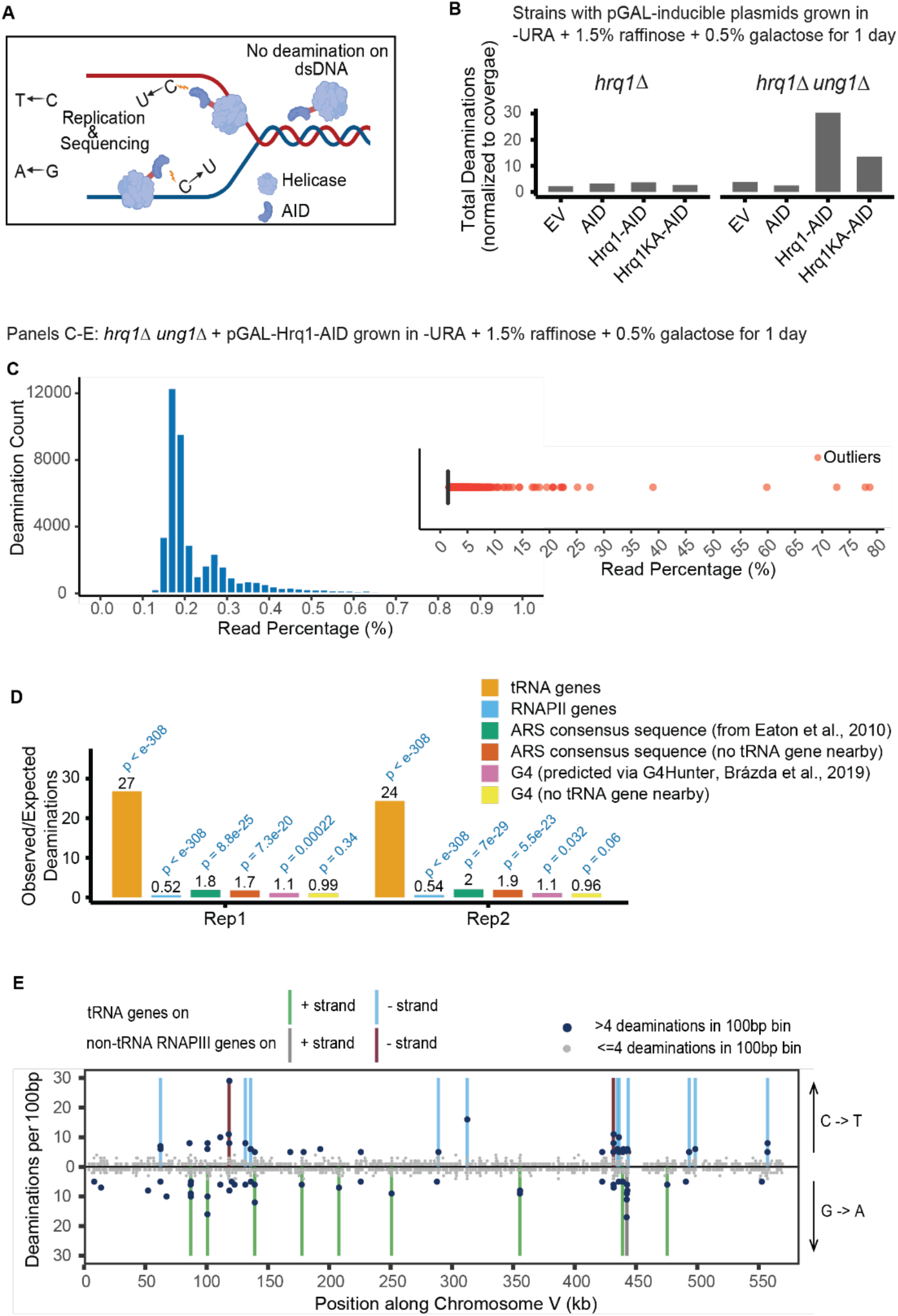
Hrq1 maps to RNAPIII-transcribed genes. **A.** Schematic for probing Hrq1 engagement with ssDNA via AID fusion. Created with BioRender.com (54). **B.** Total number of deaminations, normalized per 10⁷ mapped bases, in *hrq1Δ* and *hrq1Δ ung1Δ* strains carrying various galactose-inducible plasmids: Empty Vector (EV), AID alone, Hrq1-AID, or catalytically inactive Hrq1-K318A fused to AID (Hrq1KA-AID). **C.** Distribution of read percentages of deaminations for Hrq1-AID in the *hrq1Δ ung1Δ* strain, represented as a histogram (left; read percentage zoomed to 0-1%) and as a boxplot (right; full data range, showing high read percentage outliers as individual points). SNPs were called using a read percentage threshold of 0.1%. **D.** Ratio of observed/expected deamination at various genomic locations assessed by hypergeometric test for Hrq1-AID in the *hrq1Δ ung1Δ* strain in two independent replicate assays. Genomic locations include tRNA genes, RNAPII-transcribed genes, and ±100 bp around the midpoints of: ACS (ARS consensus sequence; from Eaton *et al.*, 2010 (43)), ACSs with no tRNA gene within this window, G4s (predicted using G4Hunter, Brázda *et al.*, 2019 (44)), and G4s with no tRNA gene within this window. *P*-values are shown above the corresponding bars. **E.** Number of Hrq1-AID-derived deaminations per 100-bp bin along a representative chromosome (ChrV) for Hrq1-AID in the *hrq1Δ ung1Δ* strain. Vertical lines show the positions of either tRNA genes or other RNAPIII-transcribed genes. Larger dots represent ≥ 5 or more deaminations in a bin, while smaller grey dots represent ≤ 4.

Genome-wide Hrq1 activity has not previously been mapped. We therefore applied our AID fusion approach in *Saccharomyces cerevisiae* to identify ssDNA regions bound by Hrq1. To minimize the accumulation of mutations during strain construction and propagation, and to generate robust deamination upon induction, we transfected a multicopy plasmid carrying *pGAL-HRQ1-AID* into *hrq1Δ* cells. We also included empty vector (EV) and plasmids with AID-only as controls, and a *pGAL-HRQ1-KA-AID* construct in which helicase-dead Hrq1 (K318A) (23) is fused to AID. Unless otherwise indicated, all strains were *ung1Δ* to prevent repair of genomic uracil via base excision repair (BER) (53). Serial dilution growth assays indicated that sustained overexpression of Hrq1-AID and Hrq1KA-AID in *ung1Δ* strains led to a modest growth defect, with Hrq1-AID more toxic than Hrq1KA-AID (Fig. S1A).

Having established these strains and their gross *in vivo* phenotypes, we next sought to identify sites of Hrq1-AID action by DNA sequencing. Following induction of Hrq1-AID expression for four doublings *via* shift to media containing 0.5% galactose, we extracted genomic DNA for Illumina sequencing. Positions where at least 0.1% of reads carried a SNP relative to the reference sequence were retained for analysis.

In *UNG1* strains, comparable numbers of deaminations were detected in Hrq1-AID and EV or AID-only controls (∼2000-5000), consistent with efficient repair of lesions by BER (Fig. 1B, S1B, Supplementary Table 2). In *ung1Δ* strains, both Hrq1-AID and the helicase-dead Hrq1KA-AID showed deamination above control levels, though Hrq1KA-AID produced fewer than half the deaminations of Hrq1-AID (EV: 5519; AID-only: 3646; Hrq1-AID: 44697; and Hrq1KA-AID: 19961) (Fig. 1B, S1B, Supplementary Table 2). Any deamination detected in the EV or AID-only controls was masked in the Hrq1-AID and Hrq1KA-AID datasets to ensure that only Hrq1-dependent deaminations were analyzed. Across the 44225 control-filtered Hrq1-AID deaminations in *ung1Δ*, the percentage of reads corresponding to C-T or G-A mutations was tightly distributed at low values (median = 0.20%, IQR = 0.18-0.28%) (Fig. 1C). A subset of deaminations (4846; 11.0% of all deaminations) exceeded the boxplot whisker threshold (Q3 + 1.5 x IQR = 0.43%), extending up to a maximum of 77.49% (Fig. 1C).

We assessed genome-wide localization using a hypergeometric observed-to-expected enrichment analysis across several genomic features, including tRNA genes, RNA polymerase II (RNAPII)-transcribed genes, ACSs (sites bound by the origin recognition complex during DNA replication), and sequences predicted to form G4 structures (Fig. 1D). To ensure ACS and G4 signals were not driven by nearby tRNA genes, ACS and G4 sequences with a tRNA gene within ±100 bp of their midpoint were excluded from a parallel analysis.

Hrq1-associated deaminations were highly enriched at tRNA genes, with 27- and 24-fold enrichment (*p* < 10^−308^) in two independent technical replicates respectively (Fig. 1D). Across the replicates, 2152 of 44225 and 1710 of 38646 unique deaminations mapped to tRNA loci, hitting 271 and 270 of 273 nuclear tRNA genes, respectively (Supplementary Table 3). All other genomic regions tested were either depleted or only mildly enriched (0.52- to 2-fold). Replication origins (ACS) showed modest but statistically significant enrichment (1.8- to 2-fold, *p* < 0.05) across both replicates; this statistical enrichment persisted when ACSs with a tRNA gene within ±100 bp of their midpoint were removed (1.7- to 1.9-fold, *p* < 0.05). G4-forming sequences were also modestly enriched (1.1-fold, *p* < 0.05) in both replicates. However, this significance was lost when tRNA-proximal G4s (±100 bp) were excluded, suggesting that the apparent association of Hrq1 with G4 motifs primarily reflects the latter’s proximity to tRNA genes.

Consistent with the statistical enrichment of Hrq1-mediated deamination at tRNA genes, analysis of 100-bp bins across the *S. cerevisiae* genome indicated that deaminations were frequently clustered close to RNAPIII-transcribed genes (Fig. 1E). C-T and G-A mutations reflect deamination on the plus or minus strand, respectively. Clusters of C-T mutations frequently coincided with RNAPIII genes coded on the minus strand, *i.e.*, those for which the plus strand is the transcriptional template. Conversely, G-A clusters co-occurred with RNAPIII genes encoded on the plus strand (transcribed from the minus strand). Of 44225 unique deaminations identified, 7454 were within ±1 kb of the transcriptional start site (TSS) of tRNA genes, with 4762 (63.9%) on the template and 2692 (36.1%) on the coding strand (binomial test, *p* = 1.46 ×10^−128^; Supplementary Table 4).

### Hrq1 engages ssDNA on the transcriptional template strand of RNAPIII genes

We quantified the deamination percentage per nucleotide, defined as the average percentage of sequencing reads with a deamination at each position across all 273 tRNA genes in the reference genome, within ±1 kb of the TSS. Deaminations peaked on the template strand, but not the coding strand, with two pronounced peaks: one at the TSS and another slightly downstream of the 3ʹ end of the tRNA (Fig. 2A). This pattern was consistent across replicates (Fig. S1C) and was not observed upon induction of AID alone (Fig. S2A), confirming that the observed pattern was due to Hrq1-driven DNA binding by AID. WRC motif frequency is comparable on the template and coding strands around tRNA genes (Fig. S2B). Normalizing the deamination percentage by the availability of potential deamination sites (WRC motif) increased the height of the downstream peak but retained strand specificity (Fig. 2B), indicating that the enrichment of deamination events on the transcriptional template strand was not due to biases in nucleotide content. We observed a similar template strand enrichment across the six non-tRNA, non-rRNA RNAPIII genes in the *S. cerevisiae* nuclear genome (*SNR6*, *RPR1*, *SCR1*, *SNR52*, *RNA170*, and *ZOD1*) with pronounced peaks upstream and within the gene body near the TSS (Fig. 2C, S1D).

**Figure 2.**
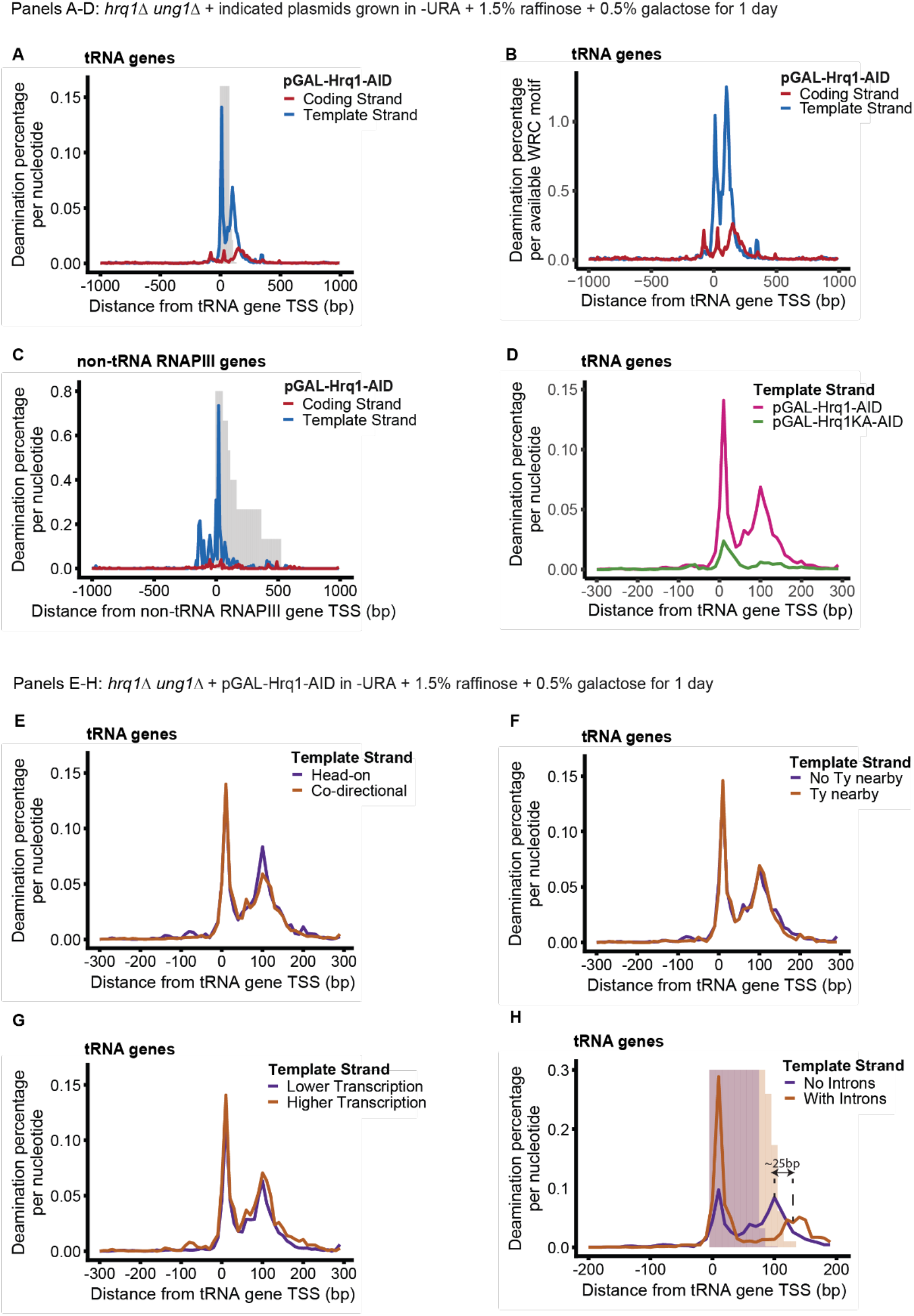
Hrq1 maps to the template strand of RNAPIII-transcribed genes. **A.** Average deamination percentage within ±1 kb of the transcription start site (TSS) across 273 tRNA genes for Hrq1-AID in *hrq1Δ ung1Δ*. Grey background bars depict a coverage histogram of the lengths of the 273 tRNAs. **B.** As for A, with deamination percentage normalized to the number of potential deamination sites (WRC or GYW depending on strand and gene orientation). **C.** As for A, for the other six non-tRNA, non-rRNA RNAPIII-transcribed genes. **D.** Average deamination percentage within ±300 bp of the TSS on the template strand of the 273 tRNA genes, comparing catalytically active Hrq1 *vs*. inactive Hrq1KA. **E-G.** Average deamination percentages within ±300 bp of the TSS on the template strand of subsets of tRNA genes binarized by: **E.** orientation relative to replication assigned by OK-seq (46), **F**. presence or absence of nearby transposable elements, and **G.** normalized transcription levels above or below the median (48). **H.** Average deamination percentage within ±200 bp of the TSS on the template strand of subsets of tRNA genes binarized by the presence or absence of introns. Background bars depict coverage histograms of the lengths of the tRNAs in the relevant subsets. Data in all panels are smoothed within 10-bp bins. Genes are oriented such that transcription proceeds left to right.

Compared to the helicase-proficient Hrq1-AID, the helicase-dead Hrq1KA-AID generated both fewer deaminations and a much weaker template-strand bias (Fig. 2D, S3A-C). Of 19533 unique deaminations in Hrq1KA-AID, 3161 mapped within ±1 kb of tRNA TSS, with approximately equal distribution between strands (1637 / 51.8% template *vs*. 1524 / 48.2% coding, binomial test *p* = 0.0463, a significant but much smaller effect size than in helicase-proficient Hrq1-AID; Supplementary Table 4). Meta-analysis of deamination frequency indicated that the 5ʹ-proximal peak was significantly reduced and the 3ʹ downstream peak largely absent (Fig. 2D, S3C). We conclude that Hrq1’s helicase activity is therefore required for its prolonged association with ssDNA on the template strand of RNAPIII genes, presumably due to DNA unwinding.

Consistent with the lack of enrichment of deamination events at specific loci outside RNAPIII-transcribed genes, we observed no substantial deviation from zero in meta-analyses of deamination frequency in regions ±1 kb of RNAPII-transcribed gene TSSs (Fig. S1E, S2C), midpoints of potential G4-forming sequences (Fig. S2D), or ACS midpoints (Fig. S2E). Average normalized coverage (per million mapped bases) across RNAPII (Fig. S2F) and tRNA genes (Fig. S2G) were roughly uniform, indicating that the observed deamination enrichment is not an artifact of sequencing coverage.

### Hrq1 is localized to nearly all tRNA genes rather than a specific subset

We next investigated whether specific subsets of tRNA genes were particularly prone to deamination by Hrq1-AID, reasoning that specific targets might point to biological mechanism. tRNAs oriented head-on *vs*. codirectionally with respect to replication-fork movement, inferred from OK-seq data (46), showed indistinguishable deamination profiles (Fig. 2E), as did tRNAs with and without nearby transposable elements (Ty) (Fig. 2F) and those with transcription levels above or below the median as determined by NET-seq (48) (Fig. 2G). Together, these data suggest that Hrq1’s presence at tRNA genes does not reflect a role in resolving transcription-replication conflicts, transposon silencing, or transcriptional regulation of specific loci.

When we compared intron-containing to intron-less tRNAs, we observed a ∼25 bp shift in the location of the downstream deamination peak, consistent with the average intron length in yeast tRNAs. The 5ʹ-proximal peak was identically positioned in both subsets, although more pronounced in intron-containing tRNAs (Fig. 2H): the increase in promoter-proximal deamination persists even when correcting for deamination motif availability (Fig. S2H). Despite this difference, the integrated template strand deamination signal was similar between the two subsets (area under the curve ratio: 1.15), consistent with similar Hrq1 activity regardless of intron status.

### Indirect fusion of AID to endogenously expressed Hrq1 v*ia* FKBP12-FRB dimerization

To exclude the possibility that Hrq1’s association with ssDNA at tRNA genes was an artifact due to overexpression, and to establish a more modular system for helicase-AID mapping, we next employed a rapamycin-inducible dimerization system to recruit AID to endogenously expressed Hrq1. We C-terminally tagged Hrq1 with FRB and expressed AID-FKBP12 from a galactose-inducible promoter. Upon galactose induction and addition of rapamycin, FKBP12 and FRB heterodimerize (55), recruiting AID to the helicase and enabling site-specific deamination at ssDNA (Fig. 3A). All experiments were performed in *ung1Δ* strains with additional *tor1-1* and *fpr1Δ* mutations to confer rapamycin resistance (49).

**Figure 3.**
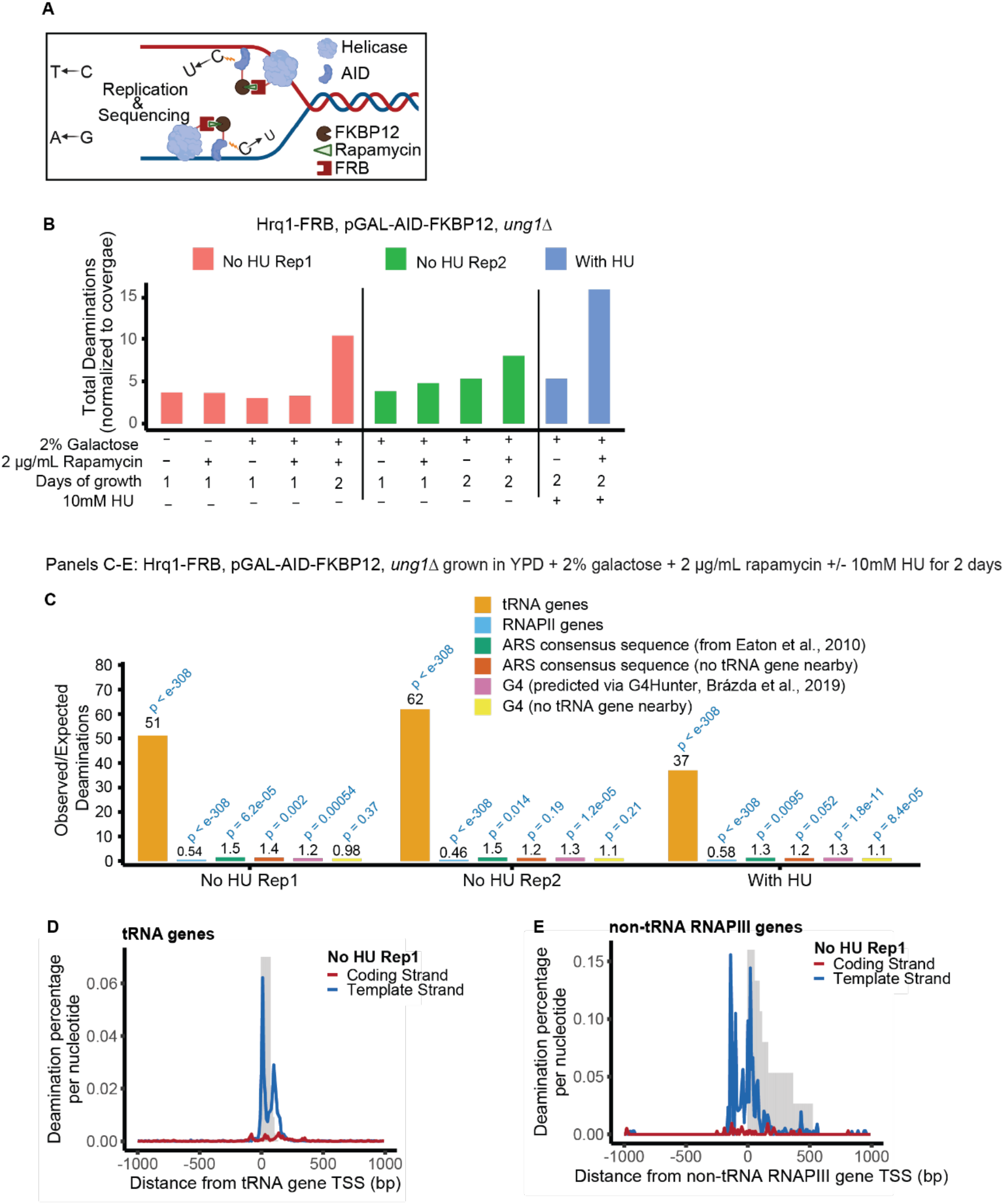
Indirect recruitment of AID to Hrq1 confirms RNAPIII-gene specificity. **A.** Schematic of the induced dimerization approach used to map ssDNA binding by endogenously expressed FRB-tagged helicases using AID. Created with BioRender.com (56). **B.** Total number of deaminations, normalized per 10⁷ mapped bases, for Hrq1-FRB-FKBP-AID under the indicated conditions. Data are shown for 1- or 2-day growth periods and without or with 10 mM HU to induce mild replication stress. **C.** Ratio of observed/expected deamination at various genomic locations assessed by hypergeometric test for Hrq1. Details as in Figure 1D. **D, E.** Average strand-specific deamination percentage within ±1 kb of the TSS across **D.** 273 tRNA genes and **E.** the other six non-tRNA, non-rRNA RNAPIII-transcribed genes. Details as in Figure 2. All strains are *ung1Δ*.

We grew *HRQ1-FRB, pGAL-AID-FKBP12 ung1Δ* cells for 5 or 10 doublings (representing 1 day of growth or 2 days with back-dilution after 1 day, respectively) in the presence of galactose and rapamycin, alongside control cultures lacking either or both, and identified deamination sites using the same pipeline as for plasmid-borne Hrq1-AID – again using a threshold of 0.1%. We detected a baseline level of deamination in control strains (5277, 5200, and 4425 in the three controls, respectively), whereas deaminations accumulated upon induction and dimerization, with higher levels observed after 2 days of growth (15100) compared to one (4865) (Fig. 3B, Supplementary Table 5). This pattern was reproducible across two biological replicates. Growth assays confirmed normal cell growth across all conditions after 2 days, with or without Ung1, indicating no growth defect or toxicity due to mutational burden (Fig. S4A). All downstream analyses therefore used the 10-doubling (2-day) condition. As done previously, deaminations identified in control conditions were excluded from experimental datasets to yield unique Hrq1-specific deaminations.

Deamination profiles across 100-bp bins after induced dimerization of Hrq1 and AID showed enrichment at RNAPIII-transcribed genes, consistent with results from Hrq1-AID overexpression (Fig. S4B; compare to Fig. 1E). We performed the same hypergeometric observed-to-expected analysis (Fig. 3C) as in the Hrq1-AID overexpression condition (Fig. 1D). tRNA enrichment was again prominent, with 51-fold and 62-fold enrichment in two biological replicates (*p* < 10^−308^) (Fig. 3C). Across the two replicates, 1353 of 14562 and 617 of 5487 unique deaminations mapped to tRNA loci, hitting 265 and 230 unique tRNA genes respectively (Supplementary Table 6). Similarly to Hrq1-AID overexpression, ACSs showed modest but statistically significant enrichment (1.5-fold, *p* < 0.05) across both replicates. However, upon removal of ACSs with a tRNA gene within ±100 bp of their midpoint, this significance was lost in one replicate but not the other. As with the Hrq1-AID strain, G4-forming sequences showed modest enrichment (1.2- to 1.3-fold, *p* < 0.05) across both replicates. Consistently, this significance was lost when tRNA-proximal G4s (±100 bp) were excluded.

Strand-specific analyses confirmed template strand enrichment at tRNA genes (Fig. 3D, S4C, Supplementary Table 7) and at the other six non-tRNA, non-rRNA RNAPIII genes (Fig. 3E), consistent with the plasmid-based expression. No enrichment was observed when AID was induced and localized to the nucleus but not dimerized with Hrq1 (Fig. S4D) or around RNAPII gene TSSs (Fig. S4E), confirming that the Hrq1 signal is RNAPIII-specific even when Hrq1 is expressed under its native promoter.

Under mild replication stress induced by 10 mM HU, induction and dimerization produced markedly higher deamination counts than the corresponding control (22312 *vs*. 9353) (Fig. 3B, Supplementary Table 5). Enrichment at tRNA genes remained pronounced at 37-fold (*p* < 10^−308^), with 1438 of 21410 unique deaminations spanning 267 tRNA genes (Fig. 3C, Supplementary Table 6), and consistent template strand enrichment (Fig. S4F, Supplementary Table 7). ACSs showed modest but statistically significant enrichment (1.3-fold, *p* < 0.05), but this significance was lost upon removal of tRNA-proximal ACSs (±100 bp). G4-forming sequences similarly showed modest enrichment (1.3-fold, *p* < 0.05), and significance was retained even after excluding tRNA-proximal G4s (±100 bp) (Fig. 3C, Supplementary Table 6).

### Hrq1 does not detectably impact RNAPIII transcription or recycling

To test whether Hrq1’s association with tRNA genes reflects a role in RNAPIII transcription, we assayed levels of the primary un-spliced Leucine tRNA transcript, pre-tL(CAA) as a proxy for RNAPIII transcriptional activity. We probed northern blots for the intron of tRNA tL(CAA) with the RNAPII-transcribed *SNR190* serving as a loading control on RNA extracted from exponentially growing wild-type (WT) and *hrq1Δ* cells at 30°C. Across four technical replicates, pre-tL(CAA) levels were comparable between WT and *hrq1Δ* strains (Fig. 4A). Thus, deletion of Hrq1 does not detectably alter RNAPIII transcription in exponentially growing cells.

**Figure 4.**
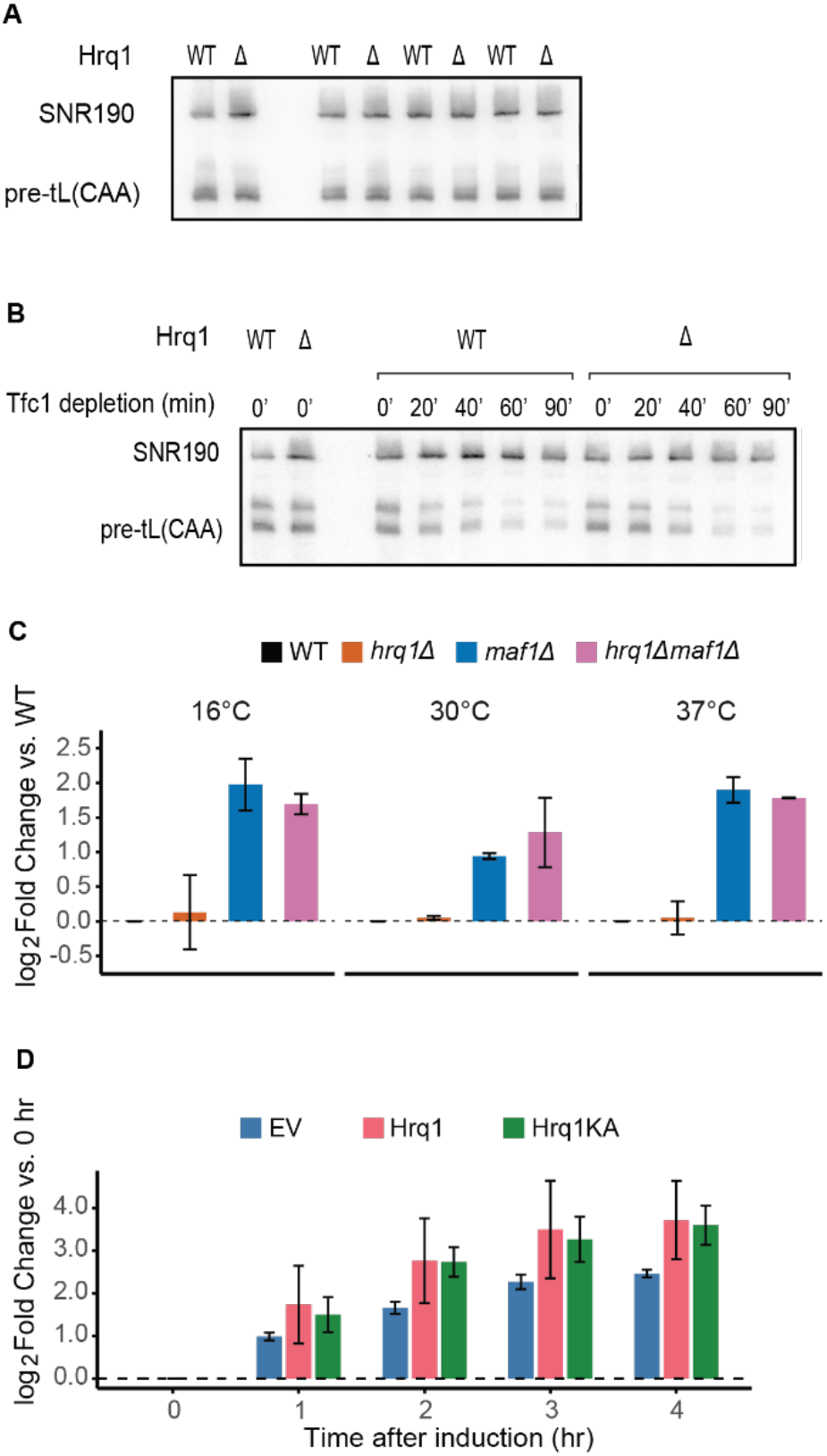
Hrq1 is not required for RNAPIII transcription or recycling. **A.** Levels of pre-tRNA tL(CAA) in wild-type (WT) *vs*. *hrq1Δ* strains across four technical replicates assessed *via* northern blotting. Northern blots were probed for the intron of the tRNA Leu(CAA), as a proxy for RNAPIII transcriptional activity. *SNR190* serves as a loading control. **B.** Pre-tL(CAA) levels over time during conditional nuclear depletion of Tfc1, a TFIIIC subunit required for RNAPIII transcription initiation. **C.** Pre-tL(CAA) levels in *hrq1Δ*, *maf1Δ*, and *hrq1Δ maf1Δ* strains grown at 16, 30, and 37°C assessed via RT-qPCR. Data are presented as log_2_-fold change relative to wild-type (WT) normalized to actin mRNA, averaged across two biological replicates. Error bars represent the standard deviation of the two replicates. Individual replicates are shown in Figure S5C. **D.** Pre-tL(CAA) levels over time following galactose-induced overexpression (2% galactose) of empty vector (EV), Hrq1, or catalytically inactive Hrq1KA in *hrq1Δ* cells, assessed via RT-qPCR at 0, 1, 2, 3, and 4 h post-induction. Data are presented as log_2_-fold change relative to EV normalized to actin mRNA, averaged across two biological replicates. Error bars represent the standard deviation of the replicates. Individual replicates are shown in Figure S5D.

To test whether Hrq1 plays a role in RNAPIII recycling, we depleted Tfc1, a TFIIIC subunit required for transcription initiation, from the nucleus *via* Anchor Away (49). pre-tL(CAA) levels declined similarly during Tfc1 depletion in exponentially growing cells in both wild-type and *hrq1Δ* strains over a 90-min time course after addition of rapamycin (Fig. 4B), ruling out an essential role for Hrq1 in RNAPIII recycling at tRNA genes. The two bands observed for pre-tL(CAA) likely correspond to distinct precursor processing intermediates but do not affect the interpretation of these results.

To validate the northern blot results and extend the analysis to additional conditions, we analyzed levels of the primary tL(CAA) transcript by RT-qPCR. Targeting the pre-spliced transcript provides a more direct readout of RNAPIII activity and avoids RT stalls at modified tRNA bases. Maf1, a negative regulator of RNAPIII transcription under both normal and stress conditions (57, 58), was included to verify assay sensitivity and to test whether Hrq1 affects tRNA transcription in the absence of this repressor. We assessed pre-tL(CAA) levels in exponentially growing wild-type, *hrq1Δ*, *maf1Δ*, and *hrq1Δ maf1Δ* cells at 16, 30, and 37°C to determine whether Hrq1 plays a role in RNAPIII transcription under temperature stress conditions (Fig. 4C). None of the strains showed notable growth differences at any temperature tested in the W303 and S288C genetic backgrounds (Fig. S5A). As expected, *maf1Δ* cells showed significantly elevated pre-tL(CAA) levels compared to wild-type at all three temperatures (average log_2_-fold change of 0.94-1.98; two-way ANOVA with Tukey post-hoc; *p* < 0.0001 at 16°C and 37°C, *p* = 0.015 at 30°C) (Fig. 4C). In contrast, *hrq1Δ* cells showed no significant difference from wild-type at any temperature tested (*p* > 0.95 at all temperatures). Furthermore, *hrq1Δ maf1Δ* double mutants were not significantly different from *maf1Δ* single mutants at any temperature (*p* > 0.5 at all temperatures), indicating that loss of Hrq1 does not further alter pre-tRNA levels even in the absence of Maf1. Results shown are averaged across two biological replicates with individual replicates shown in Fig. S5C.

To test whether overexpression of Hrq1 or catalytically inactive Hrq1KA affects pre-tRNA levels, we induced their expression from a galactose-inducible multicopy plasmid using 2% galactose in *hrq1Δ* cells in the exponential phase. Pre-tL(CAA) levels were measured by RT-qPCR at hourly intervals following induction (Fig. 4D). None of the strains showed growth defects compared to the empty vector control under inducing conditions (Fig. S5B). Pre-tL(CAA) levels increased over time following galactose induction across all conditions (Fig. 4D). However, no significant differences were observed between empty vector, Hrq1, and Hrq1KA at any timepoint (two-way ANOVA with Tukey post-hoc; *p* > 0.1 for all pairwise comparisons). As before, the data shown are averaged across two biological replicates (individual replicates in Fig. S5D). Therefore, while Hrq1 reproducibly and specifically engages ssDNA on the template strand of RNAPIII genes, it does not detectably modulate the expression of these genes under the conditions tested.

### AID recruitment maps the Rrm3 helicase to a subset of high-conflict tRNA genes

Inducible recruitment of FKBP12-tagged AID to FRB-tagged Hrq1 represents a modular strategy that can be extended to other helicases. To demonstrate feasibility beyond a single helicase, we applied an identical approach to Rrm3, a well-characterized member of the PIF1 helicase family with known functions at tRNA genes (46, 59, 60). As *S. cerevisiae* Pif1 has previously been shown to act redundantly with Rrm3 at tRNA genes (46), we analyzed Rrm3-associated deaminations both in the presence of wild-type Pif1 and in the *pif1-m2* (M40A) mutant, which lacks the nuclear isoform of Pif1 while retaining only the mitochondrial isoform. We grew *RRM3-FRB, pGAL-AID-FKBP12 ung1Δ* ± *pif1-m2* cells for 10 doublings with rapamycin and galactose or under control conditions and identified Rrm3-AID-specific deaminations as for Hrq1-AID.

Rrm3 has previously been shown to physically associate with the leading-strand polymerase Pol ε (61), and more recent structural work proposes that Rrm3 is positioned to translocate along the lagging-strand template (62). We therefore asked whether deaminations were biased towards the lagging-strand template (Fig. 5A) using replication-fork directionality from OK-seq (47). Regions in which >=75% of replication forks move in the same direction were classed as ‘strong strand bias’.

**Figure 5.**
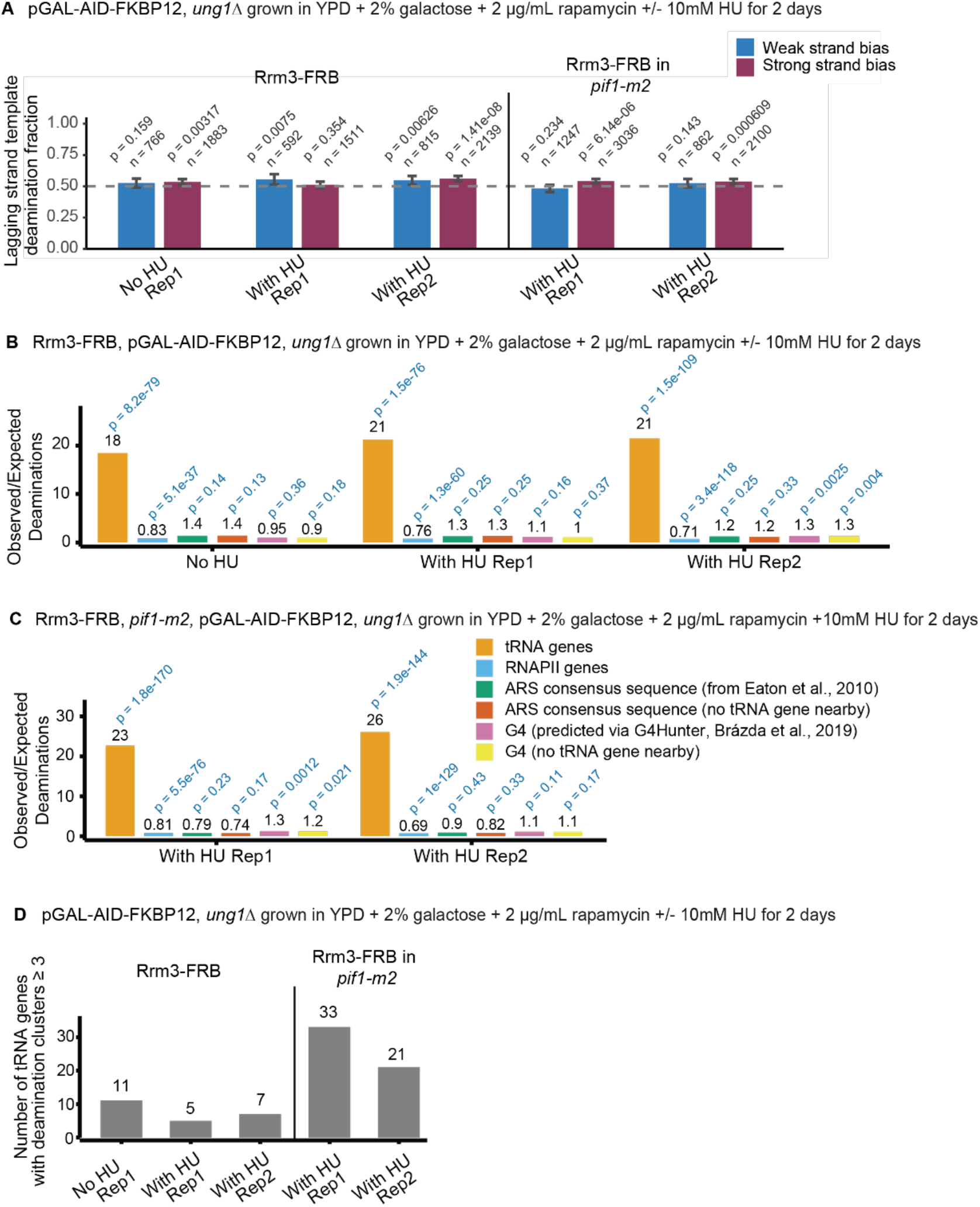
Rrm3-AID-specific deaminations are enriched at tRNA genes. **A.** Proportion of deaminations on the lagging strand template, assigned based on OK-seq data (47); 1-kb genomic windows for Rrm3 (left) and for Rrm3 in the *pif1-m2* background (right). Windows with >=75% of replication forks moving in the same direction were classed as ‘Strong strand bias’. The dashed line indicates 0.5 (no strand bias). *P*-values are from a two-sided binomial test against a null of 0.5. Error bars show 95% Clopper-Pearson confidence intervals, and n represents the number of deaminations in each subset. **B, C.** Ratio of observed/expected deamination at various loci assessed by hypergeometric test for Rrm3 in **B.** the presence and **C.** the absence of nuclear Pif1 (*pif1-m2*). Details as in Figure 1D. **D.** Number of tRNA genes that exhibit clusters of deaminations (>=3 deaminations within 100 bp) for Rrm3 (left) and for Rrm3 in the *pif1-m2* background (right). Indicated samples were grown in the presence of 10 mM HU to induce mild replication stress. All strains are *ung1Δ*.

Of 1883 deaminations in regions with a strong replication-strand bias, 1006 mapped to the lagging-strand template and 877 to the leading-strand template (proportion lagging template = 0.534, *p* = 0.00317 *via* binomial test). The remaining 766 deaminations fell within weak strand-bias windows and showed no significant strand bias (*p* = 0.159) (Fig. 5A). Neither treatment with 10 mM HU nor the absence of nuclear Pif1 substantially altered this pattern, with the proportion of deaminations on the lagging strand template ranging from 0.48 to 0.56 across all conditions and regions, and no consistent enrichment observed across replicates. Thus, while Rrm3-associated deaminations may show a slight bias toward the lagging strand template, this signal is modest in magnitude and not reproducibly detected across conditions.

We next asked whether Rrm3-associated deaminations were enriched at tRNA genes. Of 2658 unique deaminations identified, 89 overlapped with tRNA genes (Supplementary Table 8). This represents an 18-fold enrichment over expectation, (*p* < 0.05 by hypergeometric test) (Fig. 5B). Seventy-two out of 273 tRNA genes in the reference genome contained at least one deamination (Supplementary Table 8). The ratio of observed to expected deaminations at these loci was significantly higher than at other genomic features (Fig. 5B).

The enrichment at tRNA genes further increased to 21-fold under mild replication stress, with both replicates showing significant enrichment (*p* < 0.05) (Fig. 5B). Across the replicates, 81 out of 2110 and 116 out of 2979 deaminations mapped to tRNA loci, overlapping 67 and 91 unique tRNA genes, respectively (Supplementary Table 8). In contrast, tRNA gene enrichment was not observed for non-dimerized AID-FKBP12, Rrm3-FRB (Fig. S6A), or for AID dimerized with Sgs1-FRB (Fig. S6B) or Pif1-FRB (Fig. S6C), indicating that our assay does not indiscriminately mutagenize tRNA genes.

In the presence of 10 mM HU, Rrm3-associated deaminations in the *pif1-m2* background were further enriched at tRNA genes compared to *PIF1* strains, showing 23-fold and 26-fold enrichment in two biological replicates, respectively (Fig. 5C, Supplementary Table 8). Closer examination of deamination patterns revealed a subset of tRNA genes containing clusters of three or more deaminations, defined as successive deaminations occurring less than 100 bp apart (Fig. 5D). Across replicates, the number of such tRNA loci was markedly higher in the absence of nuclear Pif1 (33 and 21) compared to when nuclear Pif1 was present (11 without HU; 5 and 7 with 10 mM HU), prompting further analysis of the *pif1-m2* tRNA subset.

We tested whether the subset of tRNA genes in which Rrm3-AID induced clustered deaminations might represent those with the highest potential for transcription-replication conflicts. tRNA genes with clustered Rrm3-mediated deaminations exhibited significantly higher transcriptional activity as measured by RNAPIII occupancy (48) (Fig. 6A, *p* < 0.05 by Wilcoxon rank-sum test).

**Figure 6.**
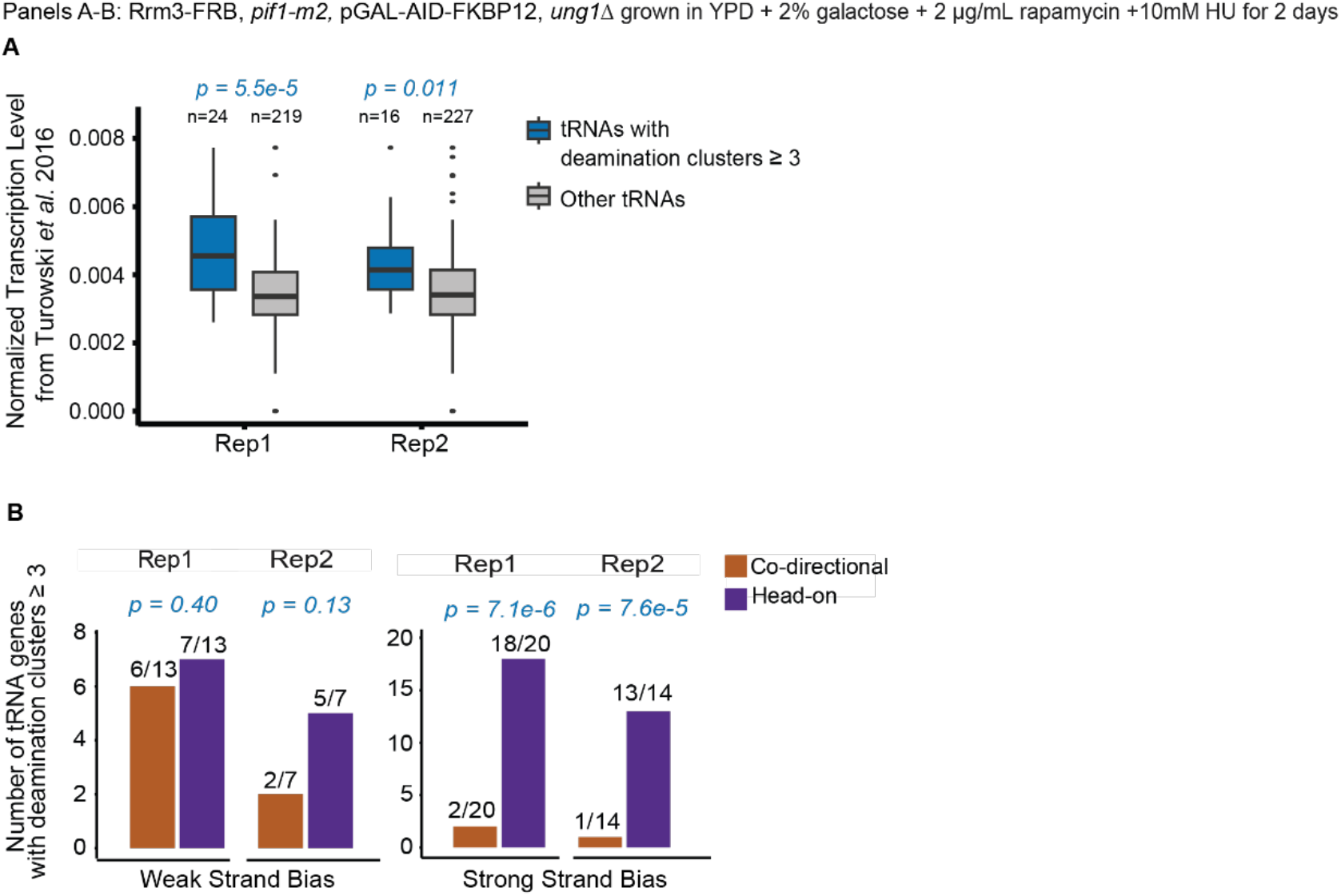
Rrm3 localizes to a subset of high-conflict tRNA genes in the absence of nuclear Pif1. **A.** Boxplots showing the normalized transcription levels of tRNAs with clustered deaminations in the absence of nuclear Pif1. *P*-values were assessed using the Wilcoxon Rank Sum Test (Mann-Whitney U) and are shown above each boxplot. Transcription levels were based on RNAPIII NET-seq data (48), and ‘n’ represents the number of tRNA genes for which data was available and included in the analysis. **B.** Orientation relative to replication of the tRNA genes that have clusters of deaminations for Rrm3 in the absence of nuclear Pif1. *P*-values from a binomial test, which accounts for the genomic proportions of head-on *vs*. codirectional tRNAs, are shown above the bar graphs. Orientation was assigned from OK-seq data (46). tRNA genes with >=75% of replication forks moving in the same direction were classed as ‘Strong Strand Bias’. Both panels A and B are for two replicate samples grown in the presence of 10 mM HU to induce mild replication stress.

Among the tRNA genes with significant clustered deaminations and strong replicative strand bias, 18 of 20 and 13 of 14 tRNA genes in two replicates were oriented head-on relative to replication (*p* < 0.05 by binomial test) (Fig. 6B). Together, these results are consistent with Rrm3’s known role in resolving transcription-replication conflicts at tRNA genes (46), confirming that AID recruitment accurately captures sites of helicase activity *in vivo*.

## Discussion

Here, we generated the first genome-wide localization data for the Hrq1 helicase in *S. cerevisiae*. Beyond its established roles in crosslink repair (23, 31, 33, 35, 37, 38, 63–65) and telomere maintenance (23–28, 33), Hrq1’s relationship to post-replication repair (PRR) has been inconsistent across organisms, variously reported as parallel to, independent of, or acting within the pathway (64–67). Given RECQL4’s links to Rothmund-Thomson syndrome, RAPADILINO, and Baller-Gerold syndrome (9–11), this equivocal functional picture, and the prior absence of genome-wide localization data for this helicase, our work addresses a gap and provides a resource for clarifying Hrq1 function.

To map helicase activity *in vivo*, we targeted AID to sites of helicase action. AID specifically deaminates cytosines on ssDNA generated by helicase unwinding, thus capturing functional helicase activity rather than mere chromatin association. To validate this approach, we applied it to Rrm3, which already has a reported genome-wide localization map. Rrm3-mediated deaminations were concentrated at tRNA genes most likely to experience transcription-replication conflicts, based on both transcriptional activity (Fig. 6A) and orientation relative to replication (Fig. 6B). This agreement with Rrm3’s known biology (46) indicates that the AID-recruitment strategy faithfully reports helicase activity *in vivo*.

Our AID-fusion approach reveals robust enrichment of Hrq1 activity across nearly all tRNA genes and other RNAPIII-transcribed loci (Fig. 1D-E), a previously uncharacterized localization. An AID-only control showed no tRNA enrichment (Fig. S2A, S4D), confirming this signal was specific to Hrq1. Deaminations occurred predominantly on the transcriptional template strand, reproducibly across Hrq1 overexpression (Fig. 2A-C) and native-promoter expression (Fig. 3B-E, S4B-F). Because Hrq1 is a 3’ to 5’ helicase, the same direction RNAPIII moves along the template strand, this strand bias suggests a functional association between Hrq1 and RNAPIII transcription machinery. Consistently, template strand enrichment was observed regardless of transcription-replication orientation, proximity to transposable elements, transcription level, or presence of introns, indicating that Hrq1 association is a general feature of RNAPIII-transcribed genes (Fig. 2E-H). Template-strand enrichment was markedly reduced in catalytically inactive Hrq1KA-AID (Fig. 2D), confirming that active helicase unwinding underlies Hrq1’s association with RNAPIII-transcribed loci. Hrq1 physically interacts with Rpb5 and, to a lesser extent, Rpb8 (32): however, these subunits are shared across all three RNA polymerases, so this interaction is unlikely to be the major driver of Hrq1’s highly specific association with RNAPIII-transcribed genes.

Despite Hrq1’s broad enrichment at RNAPIII-transcribed loci, neither deletion nor overexpression of Hrq1 affected pre-tRNA levels across multiple genetic backgrounds, growth conditions, and temperatures (Fig. 4A-D, S5A-D), indicating that Hrq1 is not required for RNAPIII transcription or recycling during exponential growth. The function of Hrq1 at RNAPIII genes therefore remains unclear. A replication-associated role is unlikely given the template-strand specificity of the signal: replication-fork passage would be expected to generate either a symmetrical deamination signal or one correlating with replication direction. A role in maintaining chromatin accessibility is similarly inconsistent with our data, as loss of Hrq1 would be expected to reduce tRNA transcription if accessibility were compromised. Instead, Hrq1’s roles in crosslink repair (23, 31, 33, 35, 37, 38, 63–65), its interaction with shared polymerase subunits Rpb5/Rpb8 (32), and its directionality matching RNAPIII suggest it may instead be recruited to resolve transcription-associated damage or R-loops – a surveillance role that would not necessarily alter steady-state pre-tRNA levels under unperturbed growth conditions if such events are rare or compensated by other pathways. Future work examining DNA damage and R-loop accumulation at tRNA genes in *hrq1Δ* may help resolve the precise function of Hrq1 at these sites.

Beyond tRNA genes, modest but significant enrichment was also observed at replication origins (ACSs) (Fig. 1D, 3C). RECQL4 is involved in replication initiation across *Xenopus*, *Drosophila*, and human cells (18, 20, 68–72) via its N-terminal Sld2-like domain (20, 68, 70), though one recent study found RECQL4 dispensable for origin firing (73). Because Hrq1 lacks this N-terminal domain and Sld2 is a separate protein in *S. cerevisiae*, any origin enrichment we observe is unlikely to reflect the same role. Consistent with this, the origin signal was modest relative to tRNA gene enrichment, and catalytically inactive Hrq1KA-AID showed similar enrichment (Fig. S3A), suggesting this signal does not reflect active helicase unwinding and is more likely an artifact of Hrq1 binding or proximity to these sites. Perhaps this is related to Hrq1’s N-terminal Cdt1-like domain (16), which may spuriously recruit Hrq1 to origins. In *S. cerevisiae* and other eukaryotes, Cdt1 is another well characterized replication initiation factor (74).

Although Hrq1 can unwind G4s *in vitro* (30), we did not detect consistent enrichment at G4s *in vivo* (Fig. 1D, 3C). G4 enrichment lost statistical significance when tRNA-proximal G4s were excluded, except in the HU-treated condition, suggesting that the apparent enrichment largely reflects proximity to tRNA loci rather than independent Hrq1 activity at G4 structures. This suggests that G4 unwinding may not be a major function of Hrq1 under normal cellular conditions or that Hrq1 may resolve G4 structures specifically at tRNAs. However, the results do not exclude activity at a subset of folded G4 structures, activity induced by replication stress, or G4 unwinding that falls below the sensitivity of the assay.

In conclusion, we present an *in vivo* sequencing-based approach that maps strand-specific cytosine deamination near a tethered protein at near-nucleotide resolution. However, the current implementation has important limitations. Deamination frequencies are low (Fig. 1C), likely reflecting AID’s relatively slow catalytic rate, which requires deep sequencing coverage and may mean that transient or rare interactions fall below the detection threshold. The low frequencies also prevent the use of Nanopore sequencing, because observed deamination frequencies are comparable to Nanopore’s error rate. Consequently, interactions with ssDNA in repetitive regions can only be mapped across aggregated repeats. Similarly, while RECQL4 has an established mitochondrial role in humans (75), the AT-rich yeast mitochondrial genome limits detection due to a paucity of cytosines. We note that tRNA genes may have abnormally high levels of ssDNA exposure due to transcription bubbles or R-loops, potentially inflating the AID signal independently of helicase unwinding and making it difficult to fully decouple transcription-derived from helicase-derived ssDNA, though the decrease in Hrq1KA-AID signal compared to Hrq1-AID indicates that helicase activity is required for maximal deamination. Future improvements, particularly the use of APOBEC family members with higher catalytic rates that retain ssDNA specificity, could increase deamination frequency, potentially enabling detection of lower-activity sites and long-read sequencing at repetitive regions. Despite these limitations, our results establish AID-fusion helicase mapping as a new tool for studying helicase biology *in vivo*, with potential for application to other ssDNA-binding proteins and organisms.

## Data Availability

Sequencing data (including custom reference genome FASTA) are available in the NCBI Sequence Read Archive under BioProject PRJNA1510710. Processed data are available at GEO accession GSE343611.

## Acknowledgements

We thank members of the Smith and Bochman labs for their valuable insights regarding this work. This work was supported in part through the NYU IT High Performance Computing resources, services, and staff expertise. We acknowledge the Zegar Family Foundation for their generous support, and thank the NYU Center for Genomics and System Biology Genomics Core for their assistance and resources.

## Author Contributions

**Shaili Regmi:** Conceptualization, Formal Analysis, Investigation, Methodology, Resources, Software, Validation, Visualization, Writing – original draft, Writing – review & editing. **Noof Alsulaiti:** Conceptualization, Formal Analysis, Funding Acquisition, Investigation, Methodology, Resources, Validation, Visualization, Writing – review & editing. **Daniel Darling:** Conceptualization, Investigation, Methodology, Writing – review & editing. **Anastasiya Bolgova:** Investigation, Resources. **Bertrand Theulot:** Investigation, Methodology, Resources, Writing – review & editing. **Spencer J. Gray:** Resources, Writing – review & editing. **Matthew L. Bochman:** Conceptualization, Formal Analysis, Funding Acquisition, Investigation, Methodology, Project Administration, Resources, Software, Validation, Writing – review & editing. **Duncan J. Smith:** Conceptualization, Formal Analysis, Funding Acquisition, Investigation, Methodology, Project Administration, Resources, Writing – original draft, Writing – review & editing.

---

**Figure S1.**
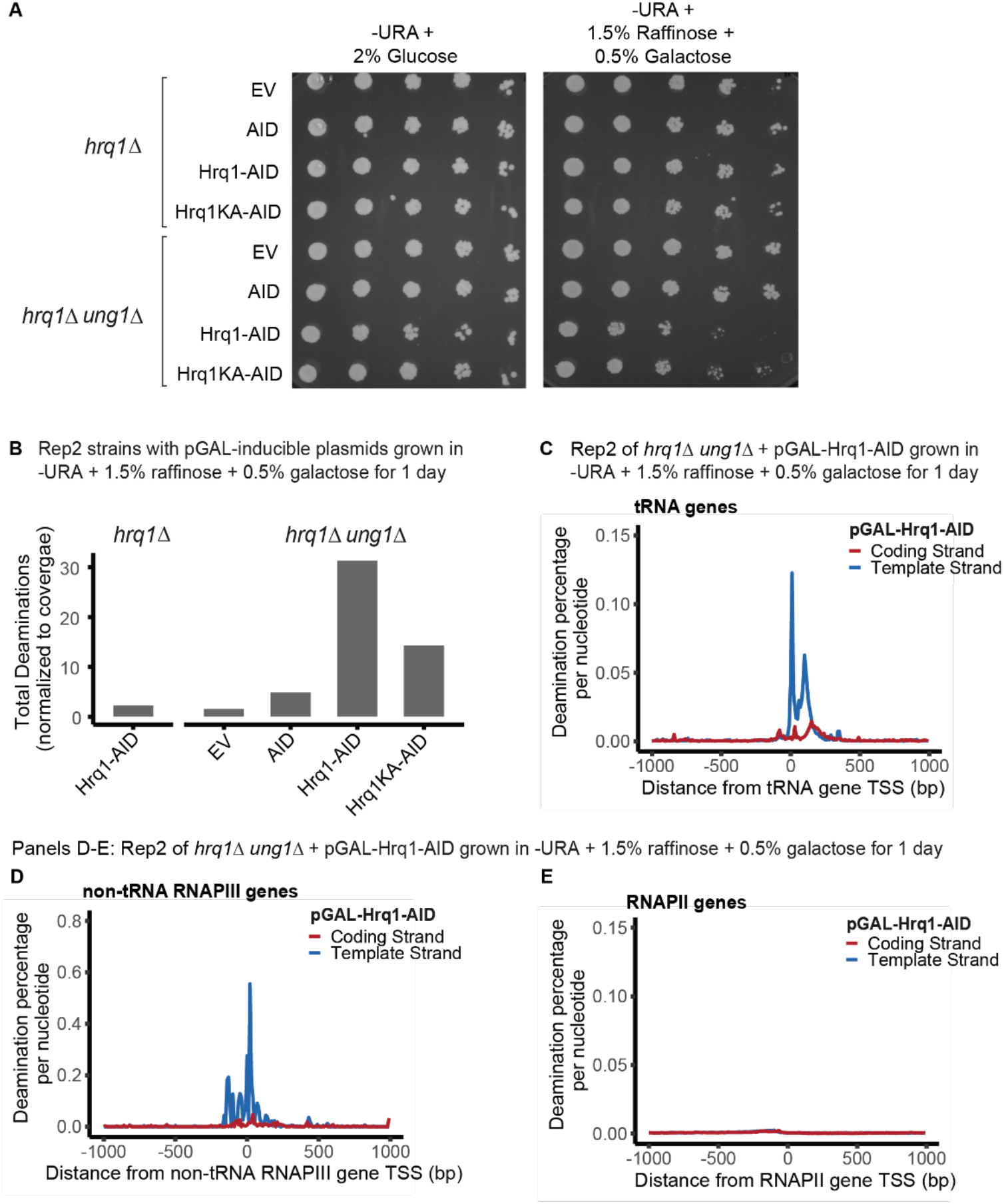
Replicates to confirm RNAPIII-specific template strand enrichment of Hrq1. **A.** Spot assays showing growth of the indicated strains with galactose-inducible plasmids detailed in Figure 1B with no induction in the presence of 2% glucose (left) and induction of the plasmids in the presence of 1.5% raffinose and 0.5% galactose (right) grown at 30°C for 2 days. Each row consists of high to low concentrations plated from left to right, respectively. **B.** Total number of deaminations, normalized per 10⁷ mapped bases, for Hrq1-AID in *hrq1Δ* and for EV, AID, Hrq1-AID, and Hrq1KA-AID in *hrq1Δ ung1Δ* from a technical replicate experiment. **C-E.** Average strand-specific deamination percentage within ±1 kb of the TSS of **C.** tRNA genes, **D.** other non-tRNA, non-rRNA RNAPIII-transcribed genes, and **E.** RNAPII-transcribed genes in the replicate assay. Details as in Figure 2.

**Figure S2.**
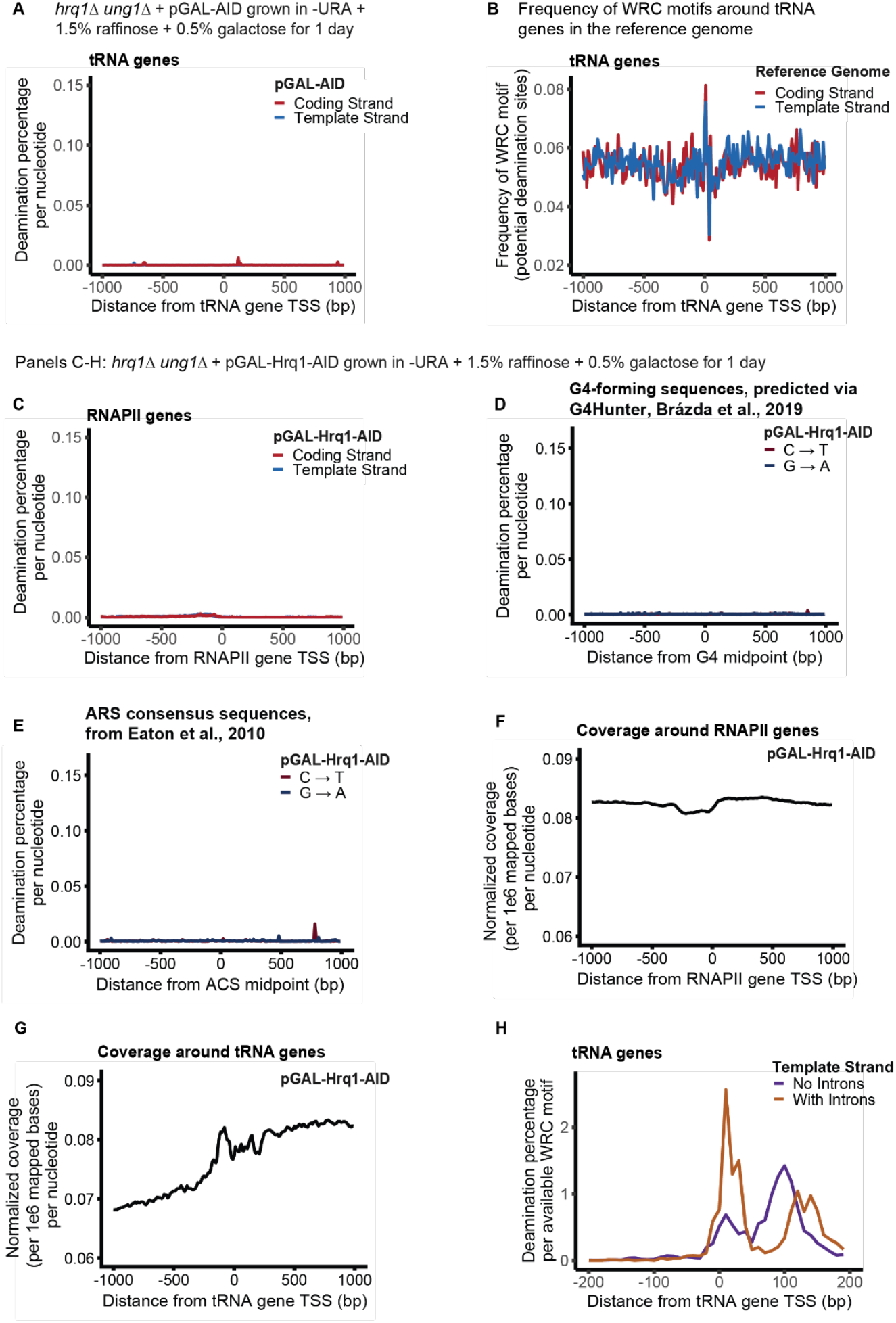
Hrq1 template-strand enrichment is specific to tRNA genes, independent of deamination motif and sequencing coverage. **A.** Average strand-specific deamination percentage within ±1 kb of the TSS across 273 tRNA genes when AID alone is induced without fusion to Hrq1 in *hrq1Δ ung1Δ*. **B.** Average strand-specific frequency of potential deamination sites (WRC or GYW depending on strand and gene orientation) in the reference genome within ±1 kb of the TSS across 273 tRNA genes. **C.** Average strand-specific deamination percentage within ±1 kb of the TSS across RNAPII-transcribed genes for Hrq1-AID. **D, E**. Average strand-specific deamination percentage within ±1 kb of the midpoint of **D.** potential G4-forming sequences (predicted using G4Hunter, Brázda *et al*., 2019 (44)) and **E.** ACSs (ARS Consensus Sequence, from Eaton *et al*., 2010 (43)). **F, G.** Average sequencing coverage (normalized per million mapped bases) after removal of duplicate and multi-mapped reads within ±1 kb of the TSS of **F.** tRNA genes and **G.** RNAPII-transcribed genes. **H.** Average deamination percentage normalized to the number of potential deamination sites within ±200 bp of TSS on the template strand of subsets of tRNA genes with or without introns. For all panels, details as in Figure 2.

**Figure S3.**
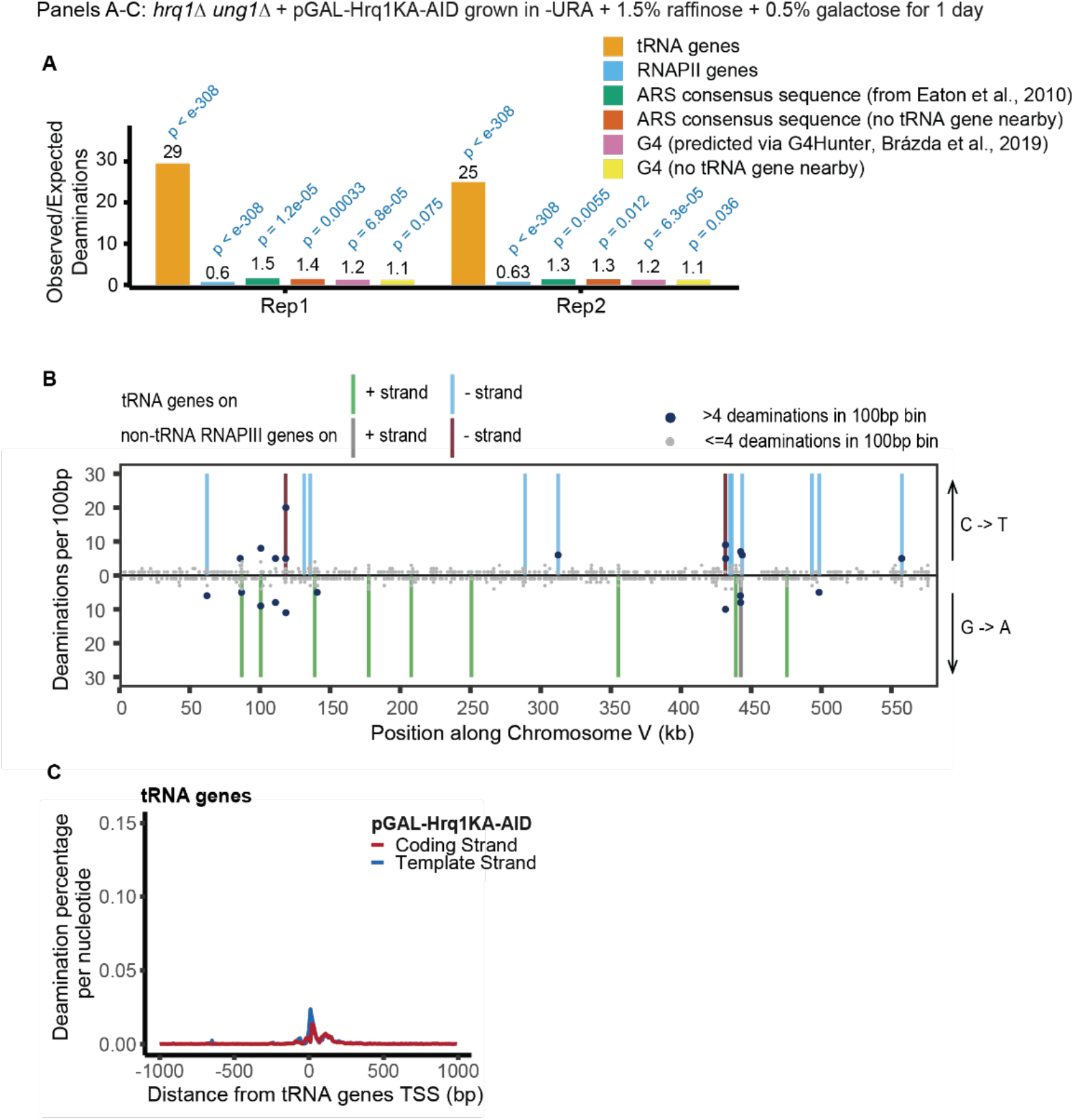
Catalytically inactive Hrq1 shows reduced enrichment at tRNA genes. **A.** Ratio of observed/expected deamination at various genomic locations assessed by hypergeometric test for Hrq1KA-AID in the *hrq1Δ ung1Δ* strain in two independent replicate assays. Details as in Figures 1D. **B.** Number of deaminations per 100-bp bin along a representative chromosome (ChrV) for Hrq1KA-AID in the *hrq1Δ ung1Δ* strain. Details as in Figure 1E. **C.** Average strand-specific deamination percentage within ±1 kb of the TSS across 273 tRNA genes for Hrq1KA-AID in *hrq1Δ ung1Δ*. Details as in Figure 2.

**Figure S4.**
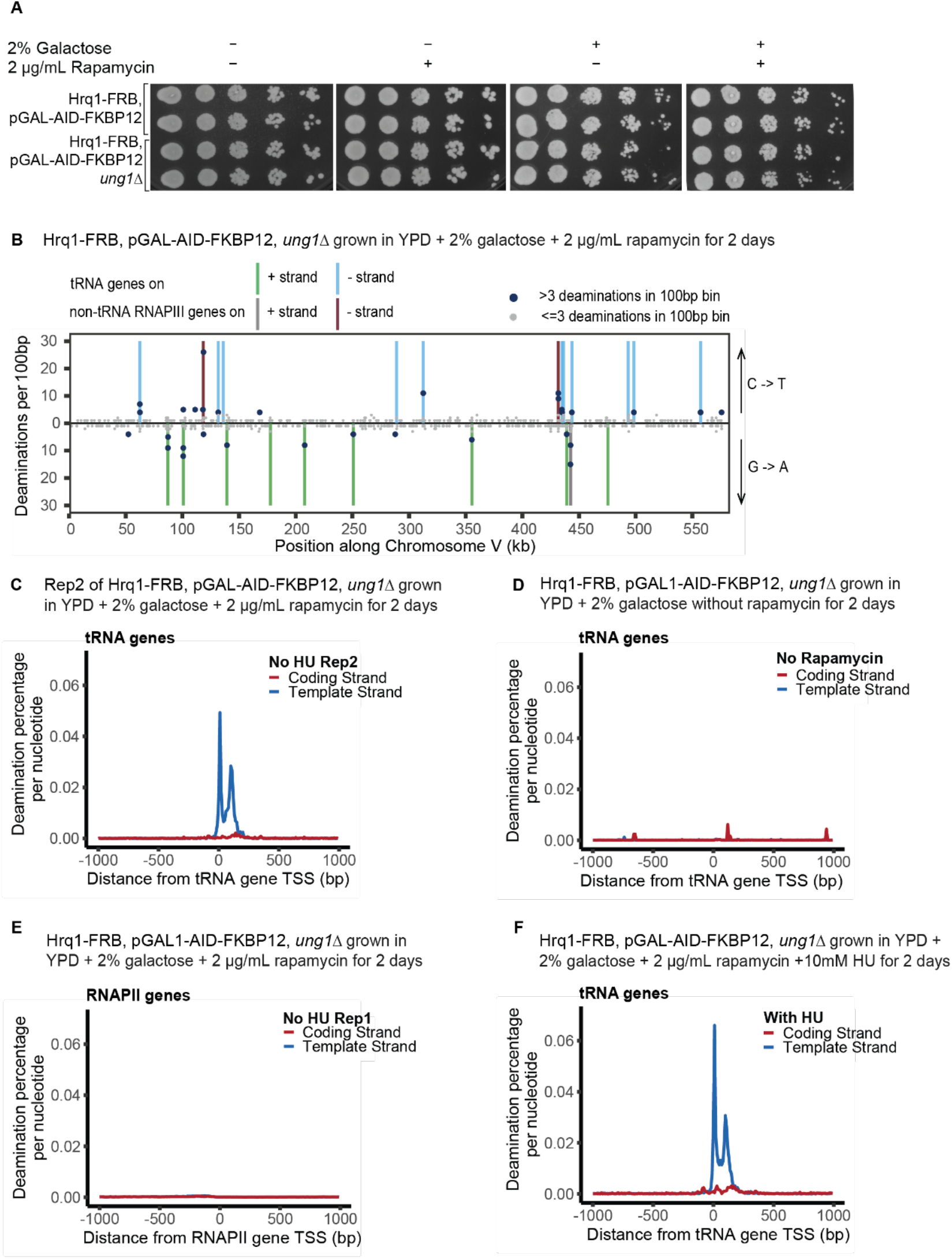
Replicates confirm endogenous Hrq1 localization at the template strand of RNAPIII-transcribed genes. **A.** Spot assays showing growth of *Hrq1-FRB, pGAL-AID-FKBP12* ± *Ung1* strains, with or without induction (2% galactose) and/or dimerization (2 µg/ml rapamycin) grown at 30°C for 2 days. Each row consists of high to low concentrations plated from left to right, respectively. **B.** Number of deaminations per 100-bp bin along a representative chromosome (ChrV). Vertical lines show the positions of either tRNA genes or other RNAPIII-transcribed genes. Larger dots represent ≥4 deaminations in a bin, while smaller grey dots represent ≤3. **C.** Average strand-specific deamination percentage within ±1 kb of the TSS across 273 tRNA genes for Hrq1 in a biological replicate strain in *ung1Δ*. **D.** Average strand-specific deamination percentage within ±1 kb of the TSS across 273 tRNA genes when AID is induced and localized to the nucleus but not dimerized with Hrq1. **E.** Average strand-specific deamination percentage within ±1 kb of the TSS across RNAPII-transcribed genes for Hrq1 in *ung1Δ*. **F.** As for C, with 10 mM HU. For panels C-F, details as in Figure 2.

**Figure S5.**
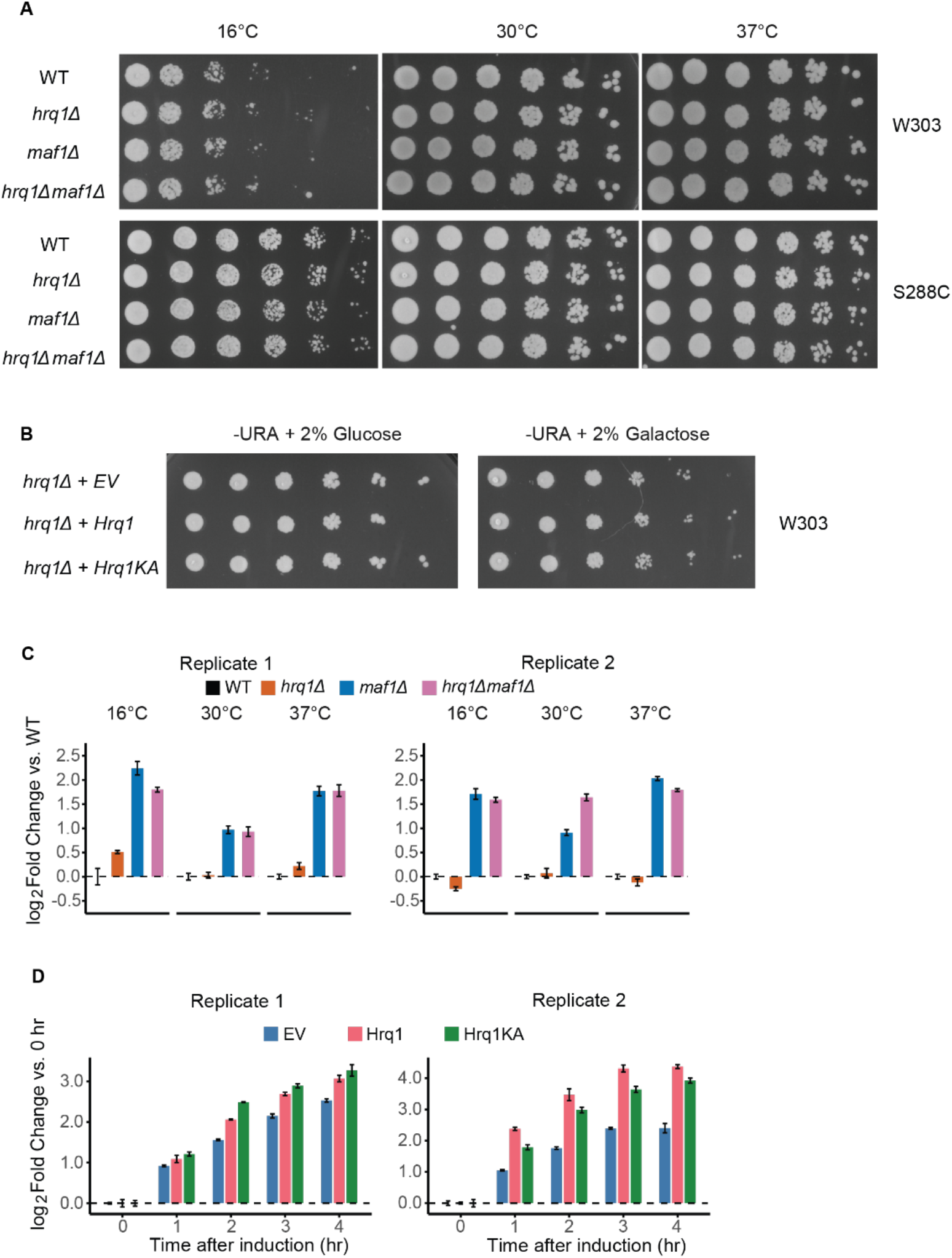
Hrq1 is not required for growth or RNAPIII transcription. **A.** Spot assays showing growth of wild-type (WT), *hrq1Δ*, *maf1Δ*, and *hrq1Δ maf1Δ* strains grown at 16°C for 8 days, and at 30°C and 37°C for 2 days in the W303 background (top) and at 16°C for 6 days, and at 30°C and 37°C for 2 days in the S288C background (bottom). Each row consists of high to low concentrations plated from left to right, respectively. **B.** Spot assays showing the growth of *hrq1Δ* cells overexpressing empty vector (EV), Hrq1, or catalytically inactive Hrq1KA under non-induced (2% glucose; left) and induced (2% galactose; right) conditions grown at 30°C for 2 days in the W303 background. Each row consists of high to low concentrations plated from left to right, respectively. **C.** Individual biological replicates for the RT-qPCR data shown in Figure 4C. Left and right panels show replicate 1 and replicate 2, respectively. Data are presented as the log_2_-fold change relative to WT normalized to actin mRNA. Error bars represent the standard deviation of three technical replicates from the same RNA sample. **D.** Individual biological replicates for the RT-qPCR data shown in Figure 4D. Left and right panels show replicate 1 and replicate 2, respectively. Data are presented as the log_2_-fold change relative to EV normalized to actin mRNA. Error bars represent the standard deviation of three technical replicates from the same RNA sample.

**Figure S6.**
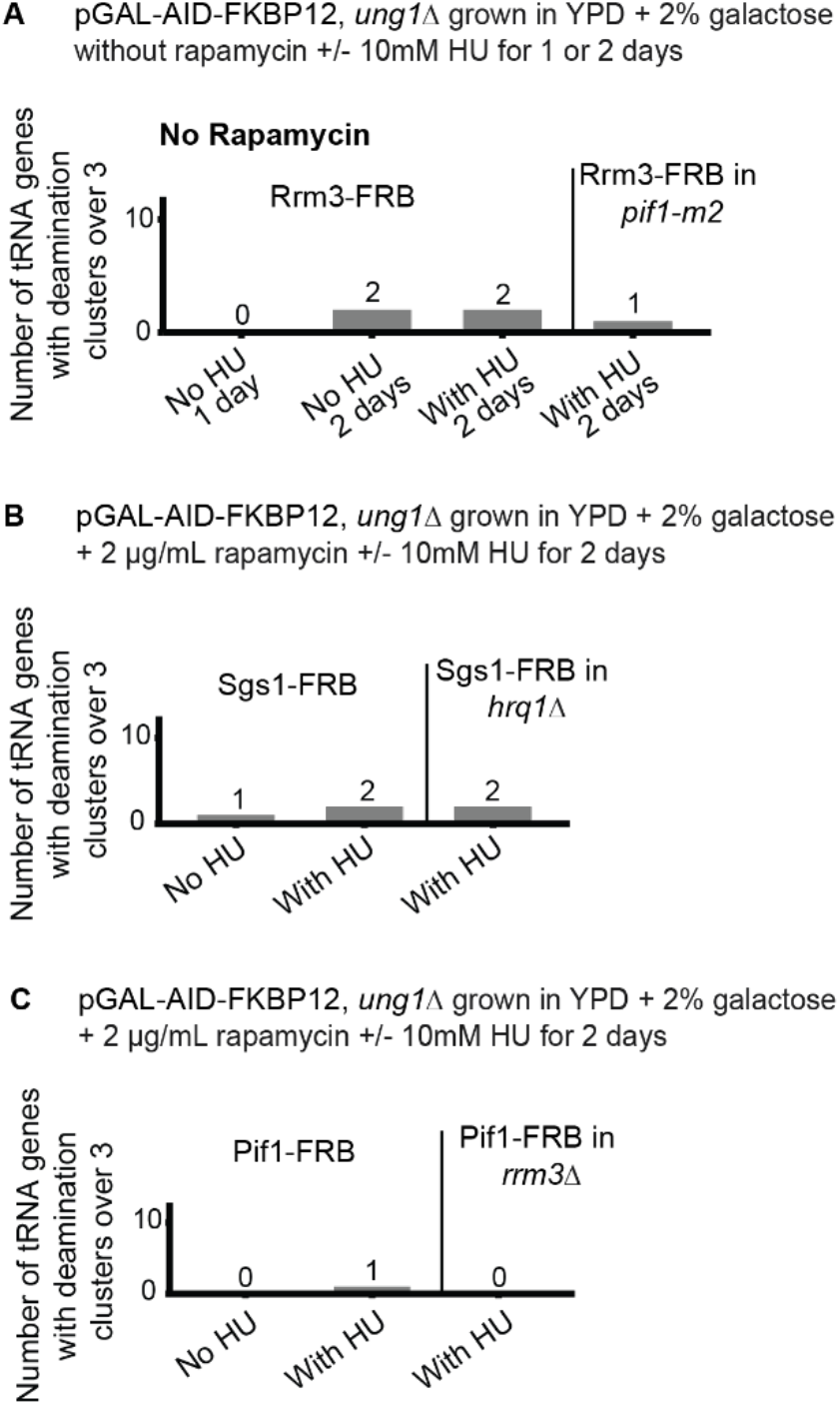
Undimerized Rrm3 and AID dimerized to other DNA-binding helicases show no tRNA gene enrichment. **A-C.** Number of tRNA genes that exhibit clusters of deaminations (>=3 deaminations within 100 bp) for **A.** AID induced and localized to the nucleus but not dimerized with Rrm3 in the presence (left) and absence (right) of nuclear Pif1, **B.** Sgs1-FRB dimerized to AID-FKBP12 in the presence (left) and absence (right) of Hrq1, and **C.** Pif1-FRB dimerized to AID-FKBP12 in the presence (left) and absence (right) of Rrm3. Indicated samples were grown in the presence of 10 mM HU to induce mild replication stress. All strains are *ung1Δ*.

