## Supplementary Tables 1-8 for "Genome-wide mapping of helicase-generated ssDNA reveals Hrq1 activity at RNA polymerase III-transcribed genes"

**Table 1.** Strains used in this study.

| **Strain** | **Genotype** | **Figure** |
| --- | --- | --- |
| W303-1A RAD5+ | *MATa ade2-1 can1-100 his3-11,15 leu2-3,112 trp1-1 ura3-1 RAD5+* | W303 MATa background / WT |
| W303-1B RAD5+ | *MATα ADE2+ can1-100 his3-11,15 leu2-3,112 trp1-1 ura3-1 RAD5+* | W303 MATα background / WT |
| S288C MATa | *MATa SUC2 gal2 mal2 mel flo1 flo8-1 hap1 ho bio1 bio6* | S288C MATa background / WT |
| S288C MATα | *MATα SUC2 gal2 mal2 mel flo1 flo8-1 hap1 ho bio1 bio6* | S288C MATα background / WT |
| yDS3331 | *W303 MATa hrq1::NAT pESC-URA (Empty Vector)* | Figure 1, S1 |
| yDS3332 | *W303 MATa hrq1::NAT pESC-URA-AID* | Figure 1, S1 |
| yDS3333 | *W303 MATa hrq1::NAT pESC-URA-Hrq1-AID* | Figure 1, S1 |
| yDS3334 | *W303 MATa hrq1::NAT pESC-URA-Hrq1-K318A-AID* | Figure 1, S1 |
| yDS3335 | *W303 MATa hrq1::NAT ung1::TRP pESC-URA (Empty Vector)* | Figure 1, 2, S1, S2, S3 |
| yDS3336 | *W303 MATa hrq1::NAT ung1::TRP pESC-URA-AID* | Figure 1, 2, S1, S2, S3 |
| yDS3337 | *W303 MATa hrq1::NAT ung1::TRP pESC-URA-Hrq1-AID* | Figure 1, 2, S1, S2 |
| yDS3338 | *W303 MATa hrq1::NAT ung1::TRP pESC-URA-Hrq1-K318A-AID* | Figure 1, 2, S1, S3 |
| yDS3361 | *W303 MATa Hrq1-V5-FRB::HYG leu2-3112::LEU2-pGal1-AID-V5-FKBP12-2xNLS ung1::TRP tor1-1::HIS3 fpr1::NatMX4* | Figure 3, S4 |
| yDS3538 | *W303 MATa Hrq1-V5-linkerV2-FRB::HYG leu2-3::LEU2-pGal1-AID-V5-linkerV2-FKBP12-2xNLS ung1::TRP tor1-1::HIS3 fpr1::NatMX4* | Figure 3, S4 |
| yDS3301 | *W303 MATa hrq1::NAT* | Figure 4, S5 |
| yDS3302 | *W303 MATa hrq1::NAT #2* | Figure 4, S5 |
| yDS3303 | *W303 MATα hrq1::NAT* | Figure 4, S5 |
| yDS3917 | *W303 MATa maf1::HYG* | Figure 4, S5 |
| yDS3918 | *W303 MATα maf1::HYG* | Figure 4, S5 |
| yDS3942 | *W303 MATa hrq1::NAT maf1::HYG* | Figure 4, S5 |
| yDS3943 | *W303 MATα hrq1::NAT maf1::HYG* | Figure 4, S5 |
| yDS777 | *W303 MATa tor1-1::HIS3 fpr1::NatMX4 RPL13A-2xFKBP12::TRP1 Tfc1-FRB::HygMX6* | Figure 4 |
| yDS3712 | *W303 MATa tor1-1::HIS3 fpr1::NatMX4 RPL13A-2xFKBP12::TRP1 Tfc1-FRB::HygMX6 hrq1::G418* | Figure 4 |
| yDS3931 | *W303 MATa hrq1::NAT pESC-URA-V5 (empty vector)* | Figure 4, S5 |
| yDS3932 | *W303 MATa hrq1::NAT pESC-URA-Hrq1-V5* | Figure 4, S5 |
| yDS3933 | *W303 MATa hrq1::NAT pESC-URA-Hrq1K318A-V5* | Figure 4, S5 |
| yDS3551 | *W303 MATa Rrm3-V5-linkerV2-FRB::HYG leu2-3::LEU2-pGal1-AID-V5-linkerV2-FKBP12-2xNLS ung1::TRP tor1-1::HIS3 fpr1::NatMX4* | Figure 5, S6 |
| yDS3552 | *W303 MATa Rrm3-V5-linkerV2-FRB::HYG leu2-3::LEU2-pGal1-AID-V5-linkerV2-FKBP12-2xNLS ung1::TRP tor1-1::HIS3 fpr1::NatMX4 #2* | Figure 5 |
| yDS3656 | *W303 MATa Rrm3-V5-linkerV2-FRB::HYG pif1-m2::G418 leu2-3112::LEU2-pGal1-AID-V5-linkerV2-FKBP12-2xNLS ung1::TRP tor1-1::HIS3 fpr1::NatMX4* | Figure 5, 6, S6 |
| yDS3657 | *W303 MATa Rrm3-V5-linkerV2-FRB::HYG pif1-m2::G418 leu2-3112::LEU2-pGal1-AID-V5-linkerV2-FKBP12-2xNLS ung1::TRP tor1-1::HIS3 fpr1::NatMX4 #2* | Figure 5, 6 |
| yDS4281 | *S288C MATa hrq1::NAT* | Figure S5 |
| yDS4283 | *S288C MATα maf1::HYG* | Figure S5 |
| yDS4296 | *S288C MATa hrq1::NAT maf1::HYG* | Figure S5 |
| yDS3545 | *W303 MATa Sgs1-V5-linkerV2-FRB::HYG leu2-3::LEU2-pGal1-AID-V5-linkerV2-FKBP12-2xNLS ung1::TRP tor1-1::HIS3 fpr1::NatMX4* | Figure S6 |
| yDS3644 | *W303 MATa Sgs1-V5-linkerV2-FRB::HYG hrq1::G418 leu2-3112::LEU2-pGal1-AID-V5-linkerV2-FKBP12-2xNLS ung1::TRP tor1-1::HIS3 fpr1::NatMX4* | Figure S6 |
| yDS3564 | *W303 MATa Pif1-V5-linkerV2-FRB::HYG leu2-3::LEU2-pGal1-AID-V5-linkerV2-FKBP12-2xNLS ung1::TRP tor1-1::HIS3 fpr1::NatMX4* | Figure S6 |
| yDS3635 | *W303 MATa Pif1-V5-linkerV2-FRB::HYG rrm3::G418 leu2-3112::LEU2-pGal1-AID-V5-linkerV2-FKBP12-2xNLS ung1::TRP tor1-1::HIS3 fpr1::NatMX4* | Figure S6 |

**Table 2.** pGAL-Hrq1-AID and controls coverage and deamination summary.

| **Dataset** | **Genotype** | **Plasmid** | **Sequencing Coverage** | **SNPs** | **Deaminations** | **C to T** | **G to A** | **% Deamination** | **Deamination Normalized to Coverage** |
| --- | --- | --- | --- | --- | --- | --- | --- | --- | --- |
| Rep1* | *hrq1Δ* | EV | 1213.06 | 139230 | 3160 | 1551 | 1609 | 2.27 | 2.170827403 |
| Rep1 | *hrq1Δ* | AID | 1202.22 | 114548 | 4674 | 2300 | 2374 | 4.08 | 3.239830434 |
| Rep1 | *hrq1Δ* | Hrq1-AID | 1211.51 | 157637 | 5337 | 2618 | 2719 | 3.39 | 3.671042598 |
| Rep1 | *hrq1Δ* | Hrq1KA-AID | 1227.92 | 195784 | 3901 | 1950 | 1951 | 1.99 | 2.647424067 |
| Rep1 | *hrq1Δ ung1Δ* | EV | 1204.73 | 138803 | 5519 | 2707 | 2812 | 3.98 | 3.817582582 |
| Rep1 | *hrq1Δ ung1Δ* | AID | 1244.3 | 152898 | 3646 | 1722 | 1924 | 2.38 | 2.441802237 |
| Rep1 | *hrq1Δ ung1Δ* | Hrq1-AID | 1227.9 | 208780 | 44697 | 22139 | 22558 | 21.41 | 30.33433099 |
| Rep1 | *hrq1Δ ung1Δ* | Hrq1KA-AID | 1233.08 | 181607 | 19961 | 9961 | 10000 | 10.99 | 13.48990462 |
| Rep2 | *hrq1Δ* | Hrq1-AID | 914.07 | 345540 | 2460 | 1181 | 1279 | 0.71 | 2.242712593 |
| Rep2 | *hrq1Δ ung1Δ* | EV | 852.1 | 173895 | 1585 | 800 | 785 | 0.91 | 1.550095932 |
| Rep2 | *hrq1Δ ung1Δ* | AID | 958.28 | 449373 | 5591 | 2663 | 2928 | 1.24 | 4.862001784 |
| Rep2 | *hrq1Δ ung1Δ* | Hrq1-AID | 1042.34 | 255897 | 39111 | 19039 | 20072 | 15.28 | 31.26847774 |
| Rep2 | *hrq1Δ ung1Δ* | Hrq1KA-AID | 953.88 | 254181 | 16314 | 8111 | 8203 | 6.42 | 14.25231685 |

*****Abbreviations used: Rep, repeat; EV, empty vector; AID, activation-induced cytidine deaminase; Hrq1KA, catalytically inactive Hrq1-K318A; SNPs, single-nucleotide polymorphisms.

**Table 3.** pGAL-Hrq1-AID in *hrq1Δ ung1Δ* deamination overlap summary.

| **Dataset** | **Total deaminations** | **Unique deaminations (control filtered)** | **Genomic region** | **Overlapping deaminations** | **Unique feature windows with >= 1 deamination** |
| --- | --- | --- | --- | --- | --- |
| Rep1 | 44697 | 44225 | tRNA | 2152 | 271 |
| Rep1 | 44697 | 44225 | RNAPII | 16168 | 4752 |
| Rep1 | 44697 | 44225 | ACS_100bp | 335 | 135 |
| Rep1 | 44697 | 44225 | ACS_no_tRNA | 314 | 133 |
| Rep1 | 44697 | 44225 | G4_100bp | 1564 | 709 |
| Rep1 | 44697 | 44225 | G4_no_tRNA | 1410 | 702 |
| Rep2 | 39111 | 38646 | tRNA | 1710 | 270 |
| Rep2 | 39111 | 38646 | RNAPII | 14556 | 4569 |
| Rep2 | 39111 | 38646 | ACS_100bp | 317 | 137 |
| Rep2 | 39111 | 38646 | ACS_no_tRNA | 295 | 135 |
| Rep2 | 39111 | 38646 | G4_100bp | 1316 | 632 |
| Rep2 | 39111 | 38646 | G4_no_tRNA | 1192 | 625 |

**Table 4.** pGAL-Hrq1-AID and pGAL-Hrq1KA-AID in *hrq1Δ ung1Δ* strand-specific tRNA deamination summary.

| **Dataset** | **Unique deaminations (control filtered)** | **Deamination withing 1 kb of tRNA TSS** | **Template strand** | **Coding strand** | **Template %** | ***P*-value (two-sided binomial)** |
| --- | --- | --- | --- | --- | --- | --- |
| Hrq1 Rep1 | 44225 | 7454 | 4762 | 2692 | 63.9 | 1.46E-128 |
| Hrq1 Rep2 | 38646 | 6284 | 4019 | 2265 | 64 | 8.81E-110 |
| Hrq1KA Rep1 | 19533 | 3161 | 1637 | 1524 | 51.8 | 0.0463 |
| Hrq1KA Rep2 | 15881 | 2288 | 1184 | 1104 | 51.7 | 0.0986 |

**Table 5.** Hrq1-FRB, pGAL-AID-FKBP12 in *ung1Δ* coverage and deamination summary.

| **Dataset** | **2% galactose** | **2** **µg/mL rapamycin** | **Days of growth** | **10 mM HU** | **Sequencing coverage** | **SNPs** | **Deaminations** | **C to T** | **G to A** | **% deamination** | **Deamination normalized to coverage** |
| --- | --- | --- | --- | --- | --- | --- | --- | --- | --- | --- | --- |
| No HU Rep1 | - | - | 1 | - | 1203.92 | 120336 | 5277 | 2663 | 2614 | 4.39 | 3.652657331 |
| No HU Rep1 | - | + | 1 | - | 1200.65 | 126010 | 5200 | 2557 | 2643 | 4.13 | 3.609144804 |
| No HU Rep1 | + | - | 1 | - | 1228.19 | 136026 | 4425 | 2176 | 2249 | 3.25 | 3.002395342 |
| No HU Rep1 | + | + | 1 | - | 1234.23 | 223890 | 4865 | 2391 | 2474 | 2.17 | 3.284770583 |
| No HU Rep1 | + | + | 2 | - | 1215.09 | 159516 | 15100 | 7361 | 7739 | 9.47 | 10.3558557 |
| No HU Rep2 | + | - | 1 | - | 1205.45 | 93310 | 5500 | 2728 | 2772 | 5.89 | 3.802176464 |
| No HU Rep2 | + | + | 1 | - | 791.02 | 50731 | 4529 | 2230 | 2299 | 8.93 | 4.771278308 |
| No HU Rep2 | + | - | 2 | - | 1018.62 | 108753 | 6436 | 3294 | 3142 | 5.92 | 5.265301062 |
| No HU Rep2 | + | + | 2 | - | 626.85 | 41417 | 6010 | 2929 | 3081 | 14.51 | 7.989745049 |
| With HU | + | - | 2 | + | 1480.43 | 217728 | 9353 | 4618 | 4735 | 4.3 | 5.264796346 |
| With HU | + | + | 2 | + | 1177.78 | 341536 | 22312 | 10906 | 11406 | 6.53 | 15.78678512 |

**Table 6.** Hrq1-FRB, pGAL-AID-FKBP12 in *ung1Δ* deamination overlap summary.

| **Dataset** | **Total deaminations** | **Unique deaminations (control filtered)** | **Genomic region** | **Overlapping deaminations** | **Unique feature windows with >= 1 deamination** |
| --- | --- | --- | --- | --- | --- |
| No HU Rep1 | 15100 | 14562 | tRNA | 1353 | 265 |
| No HU Rep1 | 15100 | 14562 | RNAPII | 5522 | 3067 |
| No HU Rep1 | 15100 | 14562 | ACS_100bp | 92 | 63 |
| No HU Rep1 | 15100 | 14562 | ACS_no_tRNA | 83 | 61 |
| No HU Rep1 | 15100 | 14562 | G4_100bp | 543 | 349 |
| No HU Rep1 | 15100 | 14562 | G4_no_tRNA | 462 | 342 |
| No HU Rep2 | 6010 | 5487 | tRNA | 617 | 230 |
| No HU Rep2 | 6010 | 5487 | RNAPII | 1755 | 1280 |
| No HU Rep2 | 6010 | 5487 | ACS_100bp | 34 | 23 |
| No HU Rep2 | 6010 | 5487 | ACS_no_tRNA | 27 | 21 |
| No HU Rep2 | 6010 | 5487 | G4_100bp | 236 | 146 |
| No HU Rep2 | 6010 | 5487 | G4_no_tRNA | 188 | 139 |
| With HU | 22312 | 21410 | tRNA | 1438 | 267 |
| With HU | 22312 | 21410 | RNAPII | 8729 | 3839 |
| With HU | 22312 | 21410 | ACS_100bp | 111 | 68 |
| With HU | 22312 | 21410 | ACS_no_tRNA | 103 | 66 |
| With HU | 22312 | 21410 | G4_100bp | 871 | 505 |
| With HU | 22312 | 21410 | G4_no_tRNA | 790 | 498 |

**Table 7.** Hrq1-FRB, pGAL-AID-FKBP12 in *ung1Δ* strand-specific tRNA deamination summary.

| **Dataset** | **Unique deaminations (control filtered)** | **Deamination within 1 kb of tRNA TSS** | **Template strand** | **Coding strand** | **Template %** | ***P*-value (two-sided binomial)** |
| --- | --- | --- | --- | --- | --- | --- |
| No HU Rep1 | 14562 | 3302 | 2556 | 746 | 77.4 | 7.64E-230 |
| No HU Rep2 | 5487 | 1532 | 1292 | 240 | 84.3 | 2.98E-174 |
| With HU | 21410 | 3858 | 2858 | 1000 | 74.1 | 1.10E-204 |

**Table 8.** Rrm3-FRB, pGAL-AID-FKBP12 in *ung1Δ* deamination overlap summary.

| **Dataset** | **Total deaminations** | **Unique deaminations (control filtered)** | **Genomic region** | **Overlapping deaminations** | **Unique feature windows with >= 1 deamination** |
| --- | --- | --- | --- | --- | --- |
| No HU | 3077 | 2658 | tRNA | 89 | 72 |
| No HU | 3077 | 2658 | RNAPII | 1545 | 1247 |
| No HU | 3077 | 2658 | ACS_100bp | 15 | 14 |
| No HU | 3077 | 2658 | ACS_no_tRNA | 15 | 14 |
| No HU | 3077 | 2658 | G4_100bp | 82 | 79 |
| No HU | 3077 | 2658 | G4_no_tRNA | 77 | 76 |
| With HU Rep1 | 2411 | 2110 | tRNA | 81 | 67 |
| With HU Rep1 | 2411 | 2110 | RNAPII | 1112 | 947 |
| With HU Rep1 | 2411 | 2110 | ACS_100bp | 11 | 11 |
| With HU Rep1 | 2411 | 2110 | ACS_no_tRNA | 11 | 11 |
| With HU Rep1 | 2411 | 2110 | G4_100bp | 77 | 74 |
| With HU Rep1 | 2411 | 2110 | G4_no_tRNA | 71 | 70 |
| With HU Rep2* | 2979 | NA | tRNA | 116 | 91 |
| With HU Rep2* | 2979 | NA | RNAPII | 1473 | 1174 |
| With HU Rep2* | 2979 | NA | ACS_100bp | 15 | 12 |
| With HU Rep2* | 2979 | NA | ACS_no_tRNA | 14 | 11 |
| With HU Rep2* | 2979 | NA | G4_100bp | 125 | 108 |
| With HU Rep2* | 2979 | NA | G4_no_tRNA | 123 | 107 |
| *pif1-m2* With HU Rep1 | 4664 | 4295 | tRNA | 177 | 113 |
| *pif1-m2* With HU Rep1 | 4664 | 4295 | RNAPII | 2423 | 1796 |
| *pif1-m2* With HU Rep1 | 4664 | 4295 | ACS_100bp | 14 | 13 |
| *pif1-m2* With HU Rep1 | 4664 | 4295 | ACS_no_tRNA | 13 | 12 |
| *pif1-m2* With HU Rep1 | 4664 | 4295 | G4_100bp | 176 | 158 |
| *pif1-m2* With HU Rep1 | 4664 | 4295 | G4_no_tRNA | 163 | 152 |
| *pif1-m2* With HU Rep2* | 2981 | NA | tRNA | 141 | 107 |
| *pif1-m2* With HU Rep2* | 2981 | NA | RNAPII | 1444 | 1165 |
| *pif1-m2* With HU Rep2* | 2981 | NA | ACS_100bp | 11 | 11 |
| *pif1-m2* With HU Rep2* | 2981 | NA | ACS_no_tRNA | 10 | 10 |
| *pif1-m2* With HU Rep2* | 2981 | NA | G4_100bp | 109 | 98 |
| *pif1-m2* With HU Rep2* | 2981 | NA | G4_no_tRNA | 106 | 95 |

* There were no galactose-only controls for these datasets.
